# Promiscuous targeting of cellular proteins by Vpr drives massive proteomic remodelling in HIV-1 infection

**DOI:** 10.1101/364067

**Authors:** Edward JD Greenwood, James C Williamson, Agata Sienkiewicz, Adi Naamati, Nicholas J Matheson, Paul J Lehner

## Abstract

HIV-1 encodes four ‘accessory proteins’ (Vif, Vpr, Vpu and Nef), dispensable for viral replication in vitro, but essential for viral pathogenesis in vivo. Well characterised cellular targets have been associated with Vif, Vpu and Nef, which counteract host restriction and promote viral replication. Conversely, whilst several substrates of Vpr have been described, their biological significance remains unclear. Here, we use complementary, unbiased mass spectrometry-based approaches to demonstrate that Vpr is both necessary and sufficient for DCAF1/DDB1/CUL4 E3 ubiquitin ligase-mediated degradation of at least 38 cellular proteins, causing systems-level changes to the cellular proteome. We therefore propose that promiscuous targeting of multiple host factors underpins complex Vpr-dependent cellular phenotypes, and validate this in the case of G2/M cell cycle arrest. Our model explains how Vpr modulates so many cell biological processes, and why the functional consequences of previously described Vpr targets, identified and studied in isolation, have proved elusive.

## Introduction

The HIV-1 ‘accessory proteins’ Vif, Vpr, Vpu and Nef function by binding host proteins and recruiting them to cellular degradation machinery, resulting in depletion of these target substrates (Matheson et al., 2016; Simon et al., 2015; Sugden et al., 2016; Sumner et al., 2017). Some of the targets of Vif, Vpu and Nef, such as Tetherin, APOBEC3 family members and SERINC3/SERINC5, are dominant acting viral restriction factors, and their degradation is therefore thought to enhance *in vivo* viral replication directly. Conversely, the role of the Vpr in enhancing viral replication remains unclear.

Unlike other accessory proteins, Vpr is packaged into nascent viral particles, delivered into newly infected cells, and therefore present in the earliest stages of the viral replication cycle. Even though Vpr does not enhance *in vitro* viral replication in most experimental systems, a number of cellular phenotypes have been ascribed to it – indeed, it has been described as an ‘enigmatic multitasker’(Guenzel et al., 2014). For example, expression of Vpr has variously been reported to cause arrest of cycling cells at the G2/M phase, apoptosis, enhancement of HIV gene expression, and stimulation or inhibition of key signalling pathways such as NFκB and NFAT(Ayyavoo et al., 1997; Felzien et al., 1998; Lahti et al., 2003; Re et al., 1995; Roux et al., 2000)

Although the mechanisms by which Vpr causes such complex effects is controversial, most reports agree that they depend on Vpr interacting with a cellular E3 ligase complex containing DCAF1, DDB1 and Cul4 (Dehart and Planelles, 2008; Le Rouzic et al., 2007). As with the other accessory proteins, Vpr is therefore presumed to function by recruiting cellular factors to this E3 ligase complex, resulting in their ubiquitin-dependent degradation. Accordingly, several host factors depleted by Vpr have been identified, but their connection to Vpr-associated cell biological phenotypes is generally unclear, as is their role in regulating viral replication in vivo (Hofmann et al., 2017; Hrecka et al., 2016; Laguette et al., 2014; Lahouassa et al., 2016; Lv et al., 2018; Maudet et al., 2013; Romani et al., 2015; Schrofelbauer et al., 2005; Zhou et al., 2016).

We previously used unbiased quantitative proteomics to map temporal changes in cellular protein abundance during HIV infection of CEM-T4 T-cells, and identify novel targets of Vpu (SNAT1), Nef (SERINC3/5) and Vif (PPP2R5A-E) (Greenwood et al., 2016; Matheson et al., 2015). Nonetheless, known accessory protein targets could only account for a tiny fraction of all the HIV-dependent protein changes observed in our experiments (Greenwood et al., 2016). Given the varied cell biological phenotypes ascribed to Vpr, we hypothesised that it may be responsible for some of the remaining changes. In this study, we therefore undertake a comprehensive analysis of the effects of Vpr on the cellular proteome of HIV-1 infected cells and combine this with further unbiased approaches to identify cellular proteins directly targeted and degraded by Vpr. Our data suggest a new model for the effects of Vpr on cells, in which promiscuous targeting of host factors distinguishes it from other HIV accessory proteins.

## Results

### Vpr is required for global proteome remodelling in HIV infected cells

First, we compared total proteomes of uninfected cells with cells infected with either WT HIV or an HIV Vpr deletion mutant (HIV ΔVpr) at an infectious MOI of 1.5 (**Figure 1A**), resulting in approximately 75% infection (**Figure 1B**). Data from this experiment are available, together with the other proteomics datasets presented here, in a readily searchable interactive format in **Table S1**. As expected, amongst the 7,774 quantitated proteins, we observed widespread changes in cells infected with wild-type HIV (**Figure 1C** left panel). Together with known Nef, Vpu and Vif, targets, we saw depletion of previously reported Vpr targets including HLTF (Hrecka et al., 2016; Lahouassa et al., 2016), ZGPAT (Maudet et al., 2013), MCM10 (Romani et al., 2015), UNG (Schrofelbauer et al., 2005), TET2 (Lv et al., 2018), MUS81 and EME1 (Laguette et al., 2014; Zhou et al., 2016). DCAF1, part of the ligase complex used by Vpr to degrade targets was also depleted, consistent with a previous report (Lapek et al., 2017).

**Figure 1.**
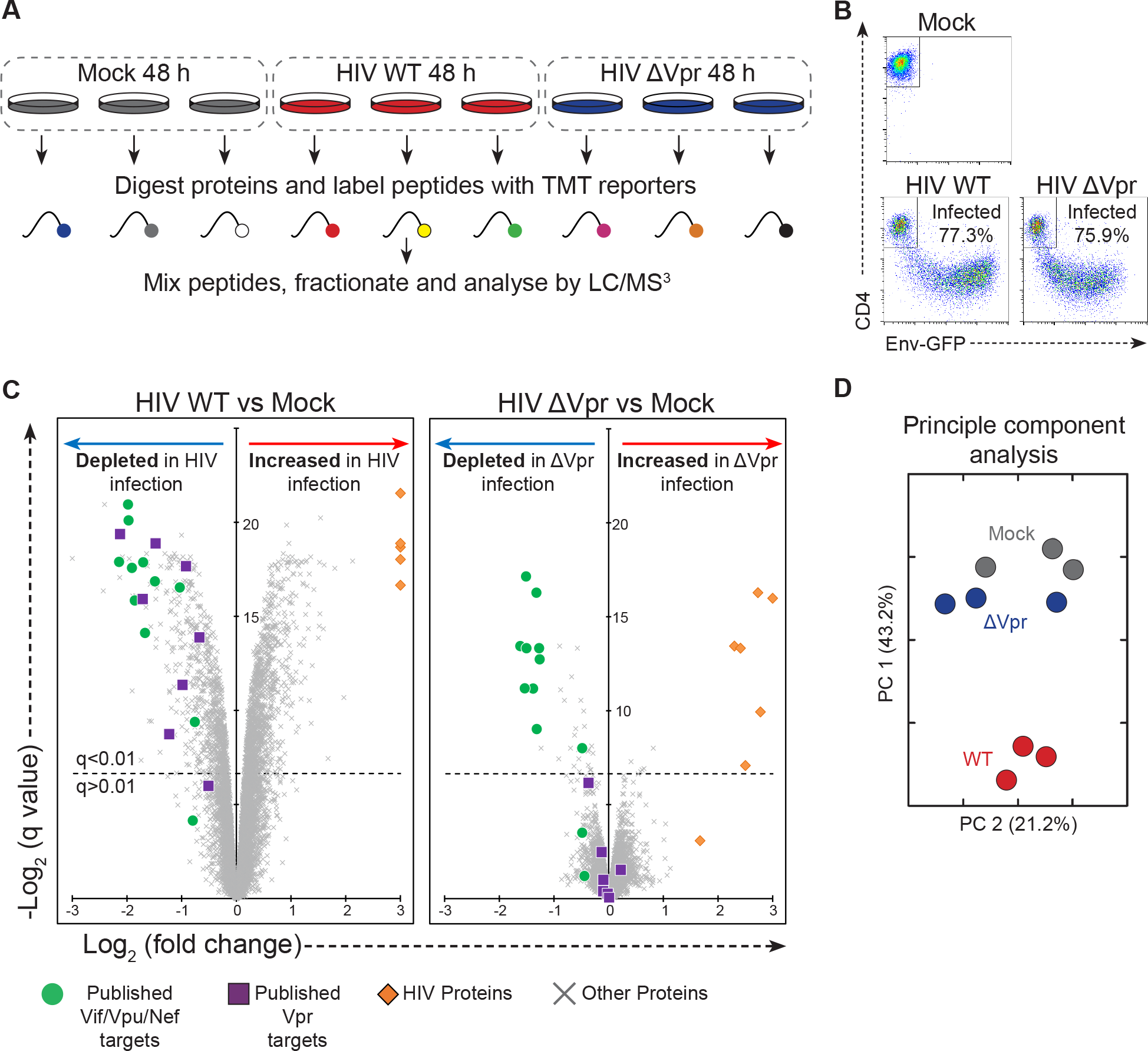
Proteomic analysis of the effect of Vpr in HIV infection. **A**, Graphical summary of the HIV and ΔVpr HIV infection TMT experiment. Three replicates of uninfected (mock), WT HIV infected and ΔVpr infected cells were prepared and analysed in parallel using TMT labelling. **B,** FACS plots showing the quantification of infection in an example replicate for each of the three conditions. Infected cells lose CD4 expression and become GFP positive. **C,** Scatterplots displaying pairwise comparisons between WT, ΔVpr and mock-infected cells. Each point represents a single protein, with HIV proteins and host proteins of interest highlighted with different symbols (see key). **D,** Principle component analysis of the samples in this experiment, with WT infected replicates in red, ΔVpr in blue and mock infected cells in grey.

In HIV ΔVpr infection (**Figure 1C**, right panel), depletion of Nef, Vpu and Vif targets was maintained. Remarkably, as well as abolishing depletion of known Vpr targets, almost all of the previously uncharacterised protein changes were also reduced or abolished in HIV ΔVpr infection. Whilst 1,944 proteins changed significantly (q<0.01) in WT HIV-infected cells, only 41 protein changes (2%) retained significance when cells were infected with HIV ΔVpr. Indeed, principle component analysis showed that cells infected with HIV ΔVpr virus share more similarity on the proteome level with uninfected cell than with cells infected with WT virus (**Figure 1D**).

### Incoming Vpr protein alone drives massive cellular proteome remodelling

Since Vpr enhances expression of other viral proteins (Forget et al., 1998; Goh et al., 1998) (**Figure 1C**), differences between WT and ΔVpr viruses could potentially be explained by secondary changes in expression levels of other proteins, or different rates of progression of WT and ΔVpr viral infections. To eliminate these potential confounders, we next examined the effect of Vpr acting alone. Unlike other HIV-1 accessory proteins, Vpr is specifically packaged into nascent viral particles. We therefore repeated our proteomic analysis using cells exposed to lentiviral particles lacking or bearing Vpr in the presence of reverse transcriptase inhibitors (RTi). This approach excludes all *de novo* viral protein expression, focussing on changes induced by incoming Vpr delivered directly by virions (**Figure 2A**). For these experiments, cells were exposed to viral particles at an infectious MOI of 0.5 (determined in the absence of RTi).

**Figure 2.**
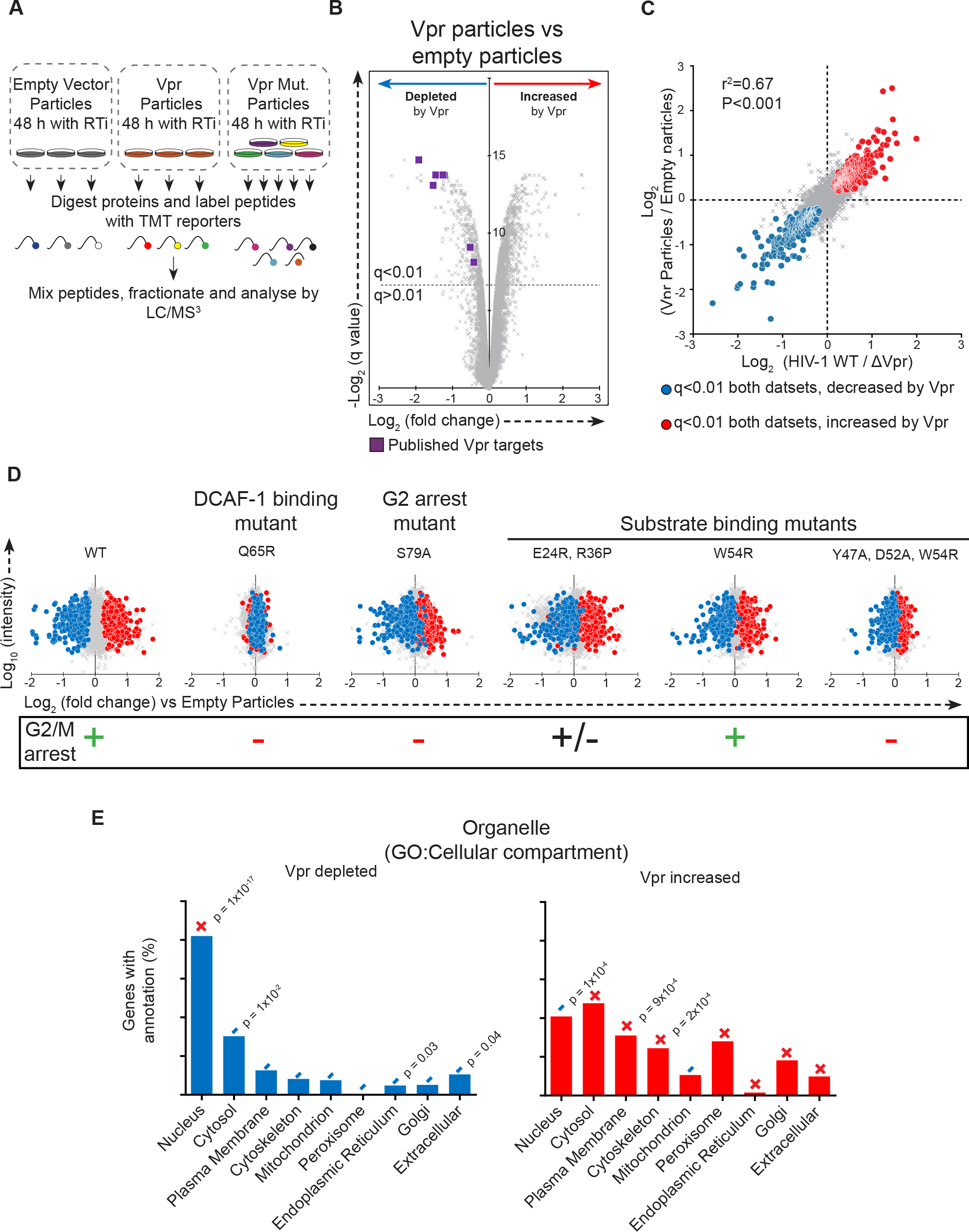
Analysis of the nature of Vpr mediated proteome remodelling. **A**, Graphical summary of the Vpr viral particle TMT experiment. Three replicates of cells exposed to empty viral particles or Vpr bearing viral particles, along with single replicates of cells exposed to viral particles bearing five different Vpr mutants were prepared and analysed in parallel using TMT labelling. **B,** Scatterplot displaying pairwise comparison between cells exposed to empty or Vpr bearing viral particles. **C**, Scatterplot comparing pairwise comparisons from two proteomics experiments, demonstrating the effect of Vpr in the context of HIV-1 infection (x-axis, as shown in **Figure 1A**) or through cellular exposure to Vpr protein alone (y-axis, as shown in **Figure 2A**). **D**, scatterplots showing the pairwise comparison between each Vpr mutant tested and empty vector control, with defined groups of 302 Vpr depleted and 413 increased proteins highlighted in blue and red respectively. **E**, Defined groups of Vpr depleted and increased proteins were subject to gene ontology enrichment analysis, compared with a background of all proteins quantitated in these experiments. GO Cellular compartment enrichment analysis results were manually curated for 9 commonly used organelle level classifications, shown here. + / − indicates if the classification was enriched or de-enriched compared with the expected number of proteins expected by chance. Where significant, associated p value represents the results of a Fisher’s exact test with Bonferroni correction. This is highly conservative as it is corrected for all 1061 possible cellular compartment terms, not just those shown.

Strikingly, changes induced by Vpr-containing viral particles phenocopied the Vpr-dependent proteome remodelling seen in HIV infection (**Figure 2B**), with a very high degree of correlation (r^2^ = 0.67; **Figure 2C**). Taking both experiments together, Vpr is therefore both necessary and sufficient to cause the significant (q<0.01) depletion of at least 302 proteins, and the upregulation of 413 – highlighted in blue and red, respectively (**Figure 2C**). This is a stringent false discovery rate and, in practice, the number of Vpr-dependent changes is almost certainly even higher. Where antibodies were available, we confirmed a proportion of these changes by immunoblot (**Figure S1A-C**).

### Cellular proteome remodelling requires interaction with DCAF1 and cellular substrates

While the function or functions of Vpr remain controversial, in all phenotypic descriptions, Vpr activity is dependent on the interaction between Vpr and the DCAF1/DDB/Cul4 ligase complex, recruitment of which results in ubiquitination and degradation of known Vpr targets by the ubiquitin-proteasome system (Dehart and Planelles, 2008). Therefore, in addition to testing the effect of wild type Vpr protein, we tested a number of previously described mutant Vpr variants (see schematic in **Figure 2A**, results in **Figures 2D & Figure S1F** and additional information in **Figure S1D-G**)

Since the Q65 residue of Vpr is required for the interaction with DCAF1 (Le Rouzic et al., 2007), we first compared proteome changes caused by a Q65R Vpr mutant with WT Vpr. As predicted, Q65R Vpr was almost completely inactive (**Figure 2D**). We recapitulated this finding by comparing the effects of WT Vpr in control cells, or cells depleted of DCAF1 (**Figure S2A**). ShRNA-mediated depletion of DCAF1 resulted in an approximately 50% reduction in protein abundance of DCAF1 (**Figure S2B**). As a proportion of cellular DCAF1 was still expressed, known Vpr effects including degradation of HLTF and upregulation of CCNB1 was partially rather than completely inhibited (**Figure S2C**). Consistent with this, Vpr mediated changes were broadly reduced in magnitude in the DCAF1 knockdown cells (**Figure S2D**). Thus, as with depletion of known Vpr targets, extensive Vpr-dependent proteomic remodelling is dependent on the interaction of Vpr with its cognate DCAF1/DDB/Cul4 ligase. Importantly, depletion of DCAF1 alone did not phenocopy Vpr-mediated proteome remodelling, and the widespread effects of Vpr are therefore unlikely to result from sequestration and/or depletion of DCAF1.

Residues E24, R36, Y47, D52 and W54 of Vpr are also required for the recruitment and degradation of previously described Vpr targets, and are reported to form the substrate-binding surface (Hrecka et al., 2016; Selig et al., 1997; Wu et al., 2016). In particular, Y47, D52 & W54 make up a proposed DNA-mimicking motif by which Vpr binds the cellular target UNG2 (Wu et al., 2016). In agreement, the Vpr_E24R, R36P_ and Vpr_W54R_ mutants showed attenuated remodelling of the proteome, while a triple mutant, Vpr_Y47A, D52A, W54R_, was defective for almost all Vpr-dependent protein changes (**Figure 2D**). Global protein remodelling therefore depends on both the substrate binding surfaces of Vpr, and the recruitment of DCAF1, suggesting that this process is mediated by recruitment of Vpr substrates to the DCAF1/DDB1/CUL4 E3 ligase complex, and their subsequent degradation.

Vpr causes G2/M arrest in cycling cells, but the mechanism remains contentious (Belzile et al., 2010; Berger et al., 2015; Fregoso and Emerman, 2016; Hohne et al., 2016; Laguette et al., 2014; Liang et al., 2015; Re et al., 1995; Romani et al., 2015; Terada and Yasuda, 2006), as is the connection to the replicative advantage Vpr provides *in vivo*. To investigate this important issue, we took advantage of previously characterised Vpr mutants. Residue S79 of Vpr is required for Vpr-dependent cell cycle arrest (Zhou and Ratner, 2000) (**Figure 2D** and **Figure S1E**). Of the other mutants we tested, Vpr_Q65R_ and Vpr_Y47A, D52A, W54R_ mutants are also unable to cause G2/M arrest, while Vpr_E24R, R36P_ has an intermediate phenotype, and Vpr_W54R_ caused G2/M arrest at wild type levels (**Figure 2D** and **Figure S1E**). Strikingly, most Vpr-dependent protein changes were also observed with the Vpr_S79A_ mutant (**Figure 2D**), and are therefore independent of G2/M cell cycle arrest. The presence or absence of G2/M arrest was also a poor correlate of proteomic remodelling across the entire panel of mutants. Cell cycle arrest therefore only explains a minority of Vpr-dependent changes **(Figure 2D)**.

To confirm this finding, we examined published datasets/reports describing proteins increased or depleted during different phases of the cell cycle, or in chemically G2/M arrested cells (Fischer et al., 2016; Ly et al., 2015) (**Figure S3**). Compared with cells exposed to Vpr in our study, cells arrested in G2 using a PLK1 inhibitor showed similar regulation of the cyclin family of proteins (**Figure S3B**), but there was little other correlation between these datasets. Thus, whilst some changes in protein levels induced by Vpr may be explained by the effects of cell cycle arrest, proteins regulated by cell cycle in these datasets only account for a small minority of Vpr-dependent changes (**Figure S3A,C,D**).

### Vpr directly targets multiple nuclear proteins with DNA/RNA-binding activity

Vpr has a nuclear localisation and all reported direct Vpr targets are nuclear proteins. Primary Vpr targets are therefore predicted to be nuclear. Conversely, secondary effects resulting from, for example, transcriptional changes, may be distributed across the cell. Analysis of the 302 proteins depleted by Vpr revealed a profound enrichment for proteins that reside in the nucleus (>80%) (**Figure 2E**). This raised the possibility that a large proportion of the proteins depleted by Vpr could be direct targets, as secondary effects should not be limited to the nucleus. Consistent with this, proteins upregulated by Vpr, which are all predicted to be secondary effects, were distributed across multiple compartments. Furthermore, proteins depleted by Vpr were enriched (>70%) for nucleic acid binding activity (**Figure S1G**). Vpr associates with DNA-binding proteins such as UNG via a substrate-binding surface that mimics DNA (Wu et al., 2016). Thus, rather than targeting a small number of cellular proteins for degradation, Vpr may have a much wider range of direct targets, and the structure of the substrate binding surface suggests a possible mechanism for promiscuous recruitment of DNA/RNA-binding cellular proteins.

To identify proteins targeted directly by Vpr, we first adopted a co-immunoprecipitation approach (**Figure S4**). Cells were transduced with a 3xHA tagged Vpr lentivirus in the presence of an shRNA to DCAF1 and the pan-cullin inhibitor MLN4924, to minimise substrate degradation and enhance co-immunoprecipitation. Factors specifically co-immunoprecipitated in the presence of Vpr are expected to include direct Vpr targets and, accordingly, were enriched for proteins depleted (rather than increased) in the presence of Vpr (**Figure S4B,D,E**). However, the co-IP was dominated by DCAF1, a stable binding partner of Vpr, identified with a signal intensity 2 orders of magnitude greater than all other proteins (**Figure S4C**). This is despite the knockdown of DCAF1 in these cells, which reduces the DCAF1 protein abundance by approximately 50% (**Figure S2B**). In addition, at least 13 proteins co-immunoprecipitating with Vpr are reported to physically interact with DCAF1 alone (Coyaud et al., 2018; Guo et al., 2016; Hossain et al., 2017), of which 11 are not regulated by Vpr, and two, CEP78 and IQGAP2, are upregulated by Vpr, explaining their presence in this list of proteins.

This mismatch between the high abundance of DCAF1 and the relatively low abundance of direct Vpr targets for degradation is consistent with previous reports, which have also found that MS-IP based techniques are ideal for the identification of the cellular machinery co-opted by viral proteins, but often struggle to identify cellular targets, which interact transiently and in competition with each other (Jager et al., 2011; Luo et al., 2016). We therefore adopted an alternative, pulsed-Stable Isotope Labelling with Amino Acids in Cell Culture (pulsed-SILAC) approach to identify host proteins specifically destabilised within 6 hrs of exposure to Vpr (**Figure 3A**). This technique is directly analogous to a traditional pulse-chase experiment using radiolabelled methionine/cysteine, but allows a global, unbiased analysis of potential cellular targets (Boisvert et al., 2012). Since proteins are fully labelled prior to exposure to Vpr, differences in abundances of labelled proteins between conditions exclusively reflect changes in protein degradation rates.

**Figure 3.**
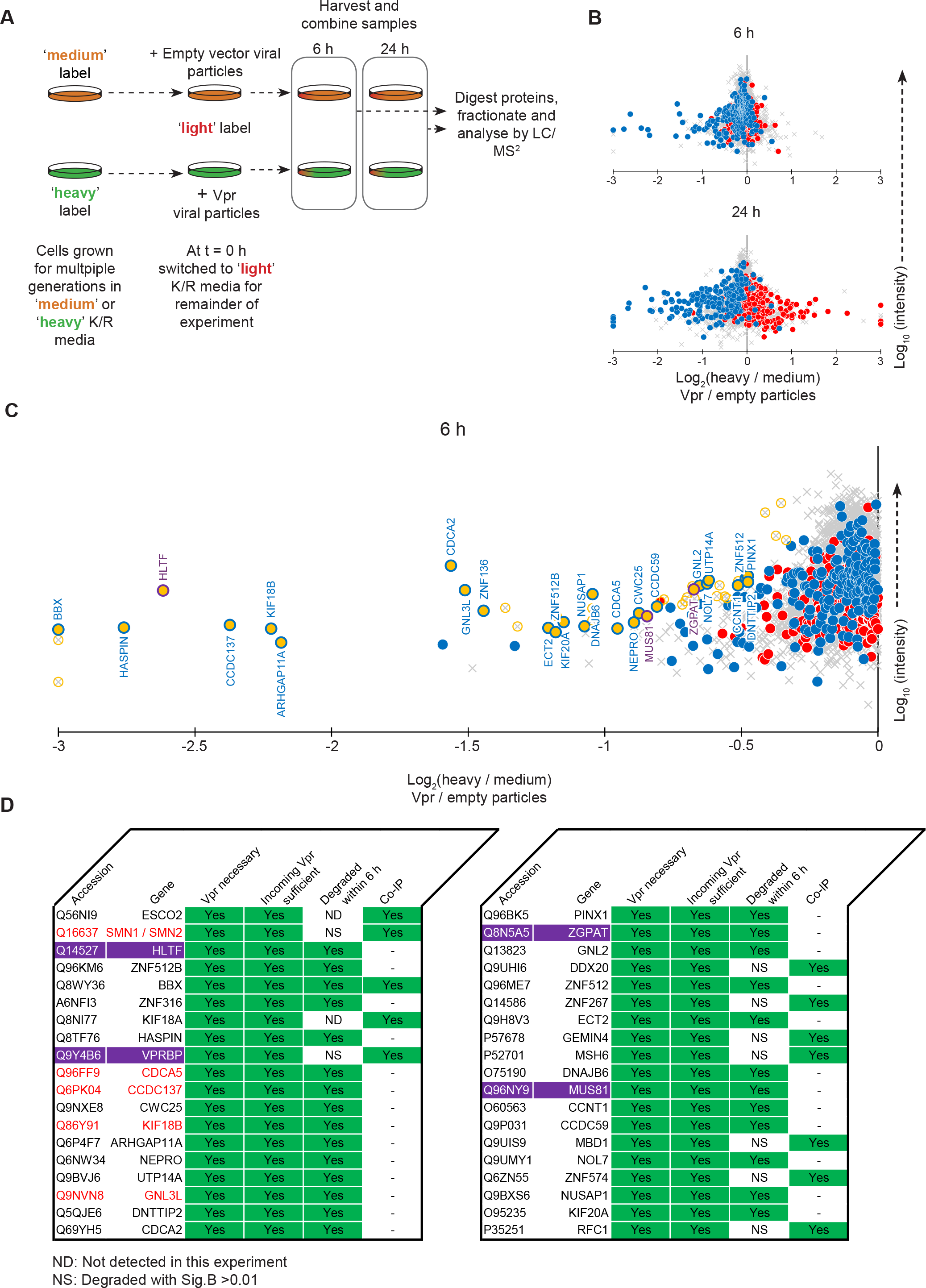
Pulsed SILAC method to identify direct targets for Vpr mediated decay. **A,** Graphical summary of the pulsed SILAC experiment. **B,** Scatterplots showing the changes to protein stability of proteins after 6 or 24 hours of exposure to Vpr bearing lentivirus compared to control lentivirus, with defined groups of 302 Vpr depleted (blue) and 413 increased (red) proteins highlighted. **C,** Expanded view of proteins degraded within 6 hours of Vpr exposure. Significantly degraded (Sig.B <0.01) proteins are highlighted in gold. Previously described Vpr targets HLTF, MUS81 and ZGPAT are shown in purple. **D,** Graphical summary of the direct targets for Vpr mediated degradation. Proteins highlighted in purple are previously described Vpr targets, proteins with red text were predicted as potential Vpr targets based on their temporal profile of depletion in Greenwood & Matheson, 2016 (Greenwood et al., 2016).

Six hours after exposure to Vpr, the stability of most proteins was unchanged (**Figure 3B,** top panel). However, a subset of proteins depleted by Vpr were already destabilised, consistent with Vpr-dependent proteasomal degradation. These 27 proteins, including HLTF, are therefore represent direct targets for Vpr-mediated depletion (**Figure 3C**). After 24 hours of exposure to Vpr, changes in protein stability reflected overall changes in protein abundance caused by Vpr in other experiments (**Figure 3B,** lower panel), including proteins with increased as well as decreased stability. These changes are therefore indicative of both direct and indirect Vpr targets.

Combining all orthogonal approaches - whole cell proteomics to identify proteins depleted by Vpr in the context of viral infection or Vpr protein alone delivered in viral particles, MS co-IP with epitope tagged Vpr, and pulsed-SILAC based identification of proteins post-translationally degraded by Vpr - we have identified at least 38 direct targets for Vpr-dependent degradation (**Figure 3D**). Vpr is both necessary and sufficient for depletion of these proteins, which are either bound by Vpr, or destabilised within 6 hours of Vpr exposure (or both). In practice, this list very likely underestimates the true number of direct Vpr targets, as several known targets of Vpr behaved appropriately, but beyond the statistical cut-offs used to derive this table (**Figure S4F**). It is also limited to proteins expressed in this T-cell model.

### Novel Vpr targets SMN1, CDCA2 and ZNF267 contribute to G2/M cell cycle arrest

Several cellular phenotypes have been described for Vpr, including G2/M arrest, transactivation of the HIV LTR, and modulation of cellular signalling pathways such as NFκB and NFAT arrest (Bolton and Lenardo, 2007; Gummuluru and Emerman, 1999; Hohne et al., 2016; Liang et al., 2015; Liu et al., 2014; Muthumani et al., 2006; Rogel et al., 1995). The mechanisms responsible for these phenotypes are controversial. Wide-scale proteome remodelling by Vpr, and direct targeting of multiple proteins, suggests a model in which Vpr interacts with diverse cellular proteins and pathways, resulting in cumulative or redundant effects on cellular phenotypes. This model does not contradict any single mechanism, but suggests that several are involved, with potential variability between different cell types and experimental systems.

To test this model, we investigated the best-described phenotype for Vpr, cell cycle arrest at the G2/M phase. We hypothesised that differential depletion of cellular proteins by different Vpr mutants tested in **Figure 2**, which displayed a spectrum of capacity to cause G2/M arrest, would highlight proteins whose depletion resulting in this cellular phenotype. We first examined proteins with a published connection to Vpr-medated G2/M arrest, MCM10, MUS81 and EME1. MCM10 is reported to be directly degraded by Vpr, resulting in cell cycle arrest (Romani et al., 2015). Vpr mediated depletion of MUS81 and EME1 was proposed to be a consequence of Vpr interaction with SLX4 (Laguette et al., 2014), another proposed mechanism of Vpr mediated G2/M arrest, though one that has been subject to conflicting reports (Berger et al., 2015; Fregoso and Emerman, 2016; Zhou et al., 2016). In our system, of these three proteins, only the depletion of MCM10 showed a strong correlation with the extent of G2/M arrest caused by the different mutants (**Figure 4A**). As previously described (Romani et al., 2015), we found RNAi depletion of MCM10 by RNAi to be sufficient to cause accumulation of cells at G2/M (**Figure 4B & 4C**).

**Figure 4.**
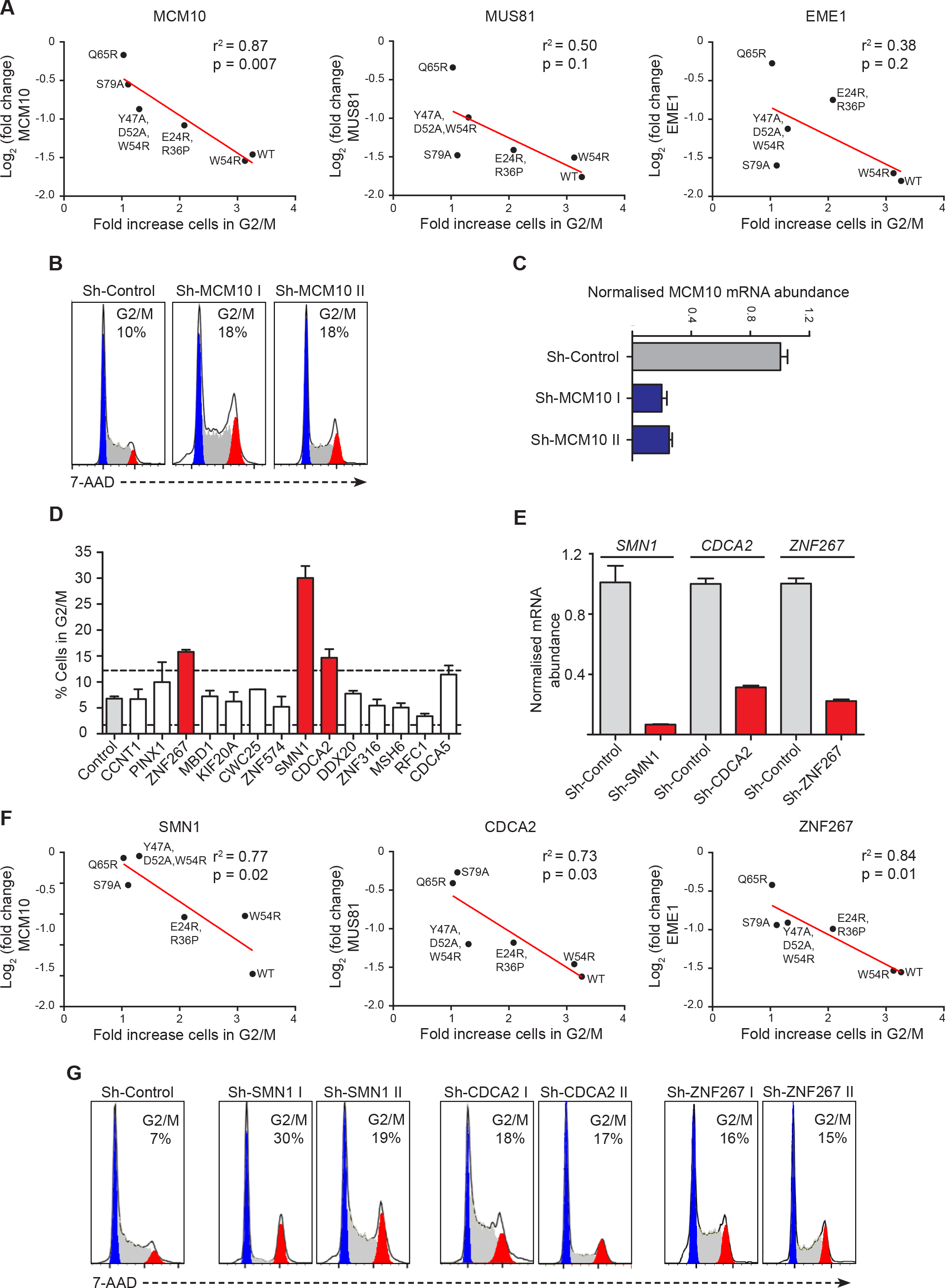
Novel Vpr targets involved in G2/M arrest. **A,** Correlation between depletion of MCM10, MUS81 and EME1 by each Vpr mutant tested in the experiment shown in **Figure 2** and the extent of G2/M arrest caused by that mutant. Red line shows linear regression analysis. **B,** Example DNA staining showing G2/M arrest caused by shRNA mediated depletion of MCM10, representative of three independent experiments. Watson pragmatic modelling was used to identify cells in G1 (blue), S (grey) or G2/M (red) phase. **C**, Real-time qRT-PCR analysis of MCM10 mRNA abundance in cells transduced with control or MCM10 targeting shRNA. Values were generated using the ΔΔCT method, relative to GAPDH mRNA abundance and normalised to the control condition. Bars show mean values, error bars show SEM of three technical replicates. **D,** Targeted shRNA screen of direct Vpr target proteins identified here, whose depletion correlated with G2/M arrest in **Figure 2**. Bars show averages of at least two replicates from more than three independent experiments, error bars showing SEM. Dashed lines show control average +/− 3 standard deviations. Control condition contains combined data from three different control shRNA. **B,** Real-time qRT-PCR analysis of mRNA abundance in cells transduced with control or targeting shRNA. Values were generated using the ΔΔCT method, relative to GAPDH mRNA abundance and normalised to the control condition. Bars show mean values, error bars show SEM of three technical replicates. **C,** Example DNA staining showing G2/M arrest caused by shRNA mediated depletion of SMN1, CDCA2 and ZNF267, representative of at least two independent experiments.

We therefore interrogated our Vpr mutant dataset (**Figure 2**) for other direct Vpr targets (**Table 1)** which, like MCM10, correlated with the extent of G2/M arrest. In total, we identified 14 targets with a significant relationship (p < 0.05 in a linear regression analysis) **(Figure 4D)**. Next, we tested whether shRNA-mediated depletion could phenocopy Vpr-dependent cell cycle arrest at G2/M (**Figure 4D)**. Depletion of 3 Vpr targets: SMN1, CDCA2 and ZNF267 caused G2/M arrest (**Figure 4D&E**), and these phenotypes were confirmed with a second shRNA (**Figure 4G**). Thus, several Vpr targets contribute independently to G2/M cell cycle arrest, consistent with a model whereby depletion of multiple cellular proteins underpin the various phenotypes associated with Vpr expression.

### Some Vpr targets are conserved across primate lentiviruses, but massive cellular proteome remodelling is unique to the HIV-1/SIVcpz lineage

Targeting of key cellular proteins such as BST2 or the APOBEC3 family is conserved across multiple lentiviral lineages, demonstrating the *in vivo* selective advantage of these interactions. We therefore tested a diverse panel of lentiviral Vpr proteins to determine if they shared activity with the NL4-3 Vpr variant used in all the experiments above. We included Vpr variants from primary isolates of HIV-1 from two distinct cross-species transmissions from apes to humans (Group M and Group O), in addition to a closely related SIVcpz variant. We also tested Vpr variants from divergent primate lineages, including HIV-2, SIVsmm, SIVagm and SIVrcm (**Figure 5A** and **Figure S5A&S5B**). In addition to Vpr, which is present in all primate lentiviruses, viruses of some lineages also bear Vpx, a gene duplication of Vpr. Since depletion of some substrates and cellular functions switches between Vpr and Vpx in lineages encoding this accessory gene (Fletcher et al., 1996; Lim et al., 2012), we also included a Vpx variant from HIV-2.

**Figure 5.**
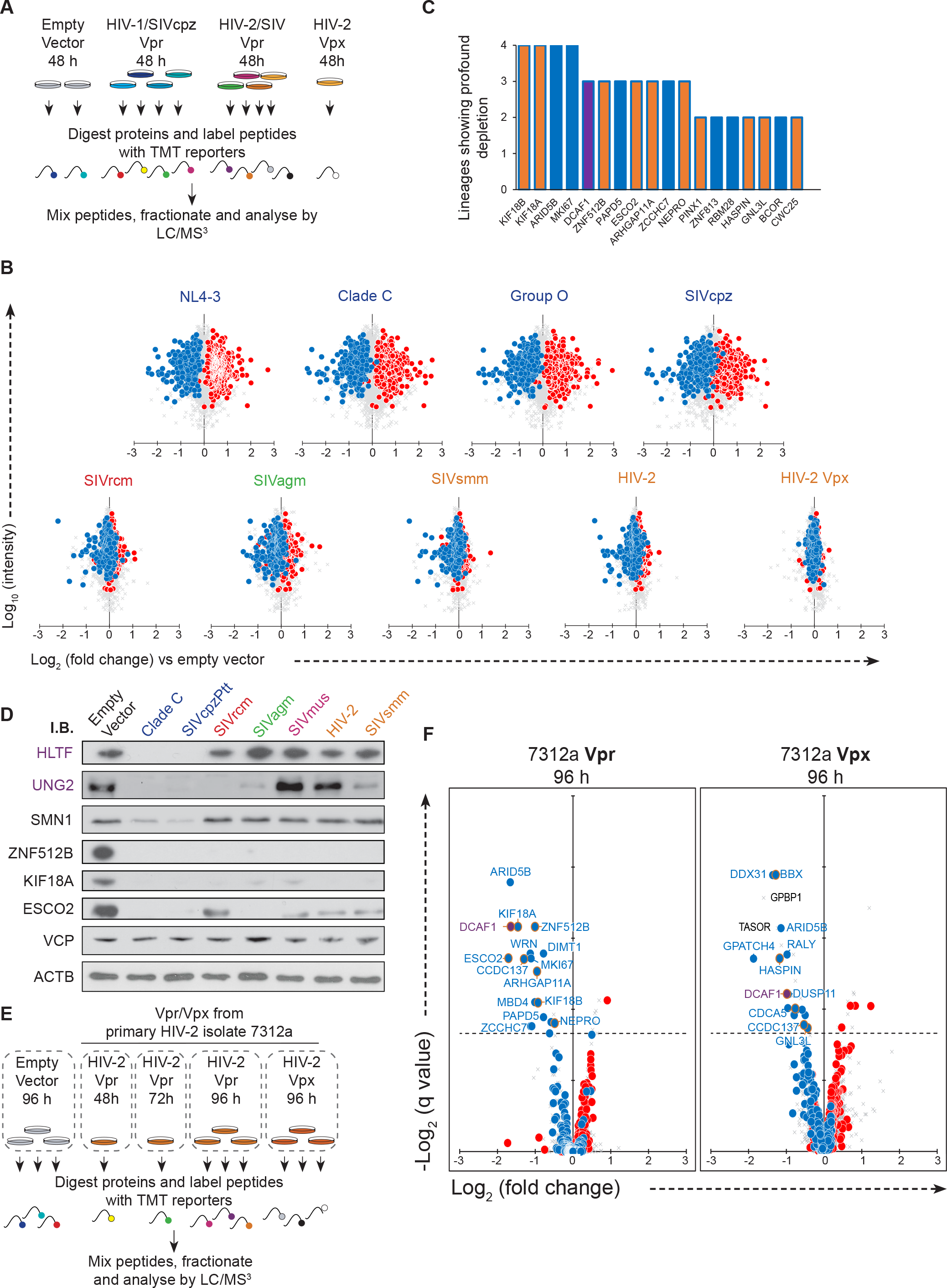
Identification of proteome changes conserved between human and primate lentiviral Vpr lineages. **A**, Graphical summary of the TMT experiment testing conservation of Vpr functions. **B**, Scatterplots showing the pairwise comparison between each Vpr tested and empty vector control with defined groups of 302 Vpr depleted (blue) and 413 increased (red) proteins highlighted. **C,** Bar chart showing proteins from the significantly Vpr depleted group that were profoundly (more than 50%) depleted in more than one lineage, of the four lineages tested (HIV-1/SIVcpz, HIV-2/SIVsmm, SIVagm, SIVrcm). Gold bars indicate proteins identified as direct targets for Vpr mediated degradation, DCAF1, known to interact with Vpr from multiple lineages, is highlighted in purple. **D**, Immunoblot of example known, non-conserved and conserved targets of Vpr mediated depletion. N.B. The HIV-2 Vpr is a primary isolate HIV-2 Vpr (7312a), while the proteomics experiment described in Figure 5A included HIV-2 ROD Vpr. **E**, Graphical summary of the TMT experiment to examine proteome changes in cells transduced with HIV-2 Vpr and Vpx for an extended period. **F,** Scatterplots displaying pairwise comparison between cells transduced with 7312a HIV-2 Vpr and Vpx for 96 h compared with those transduced with empty vector for 96 h. Blue and red dots represent the defined groups or proteins depleted or increased by NL4-3 Vpr respectively. Points ringed in gold indicate direct targets for NL4-3 Vpr mediated degradation listed in Table 1.

Extensive Vpr-dependent remodelling of the cellular proteome was conserved across the HIV-1/SIVcpz lineage (**Figure 5B** – top row). Whilst Vpr variants from other lineages showed a narrower set of changes, depletion of some proteins, particularly those most heavily depleted by HIV-1, was conserved by Vpr variants across multiple lineages (**Figure 5B,C**). Depletion of selected proteins for which commercial antibody reagents were available was readily confirmed by immunoblot of cells transduced with an overlapping panel of Vpr variants (**Figure 5D**). These conserved targets of direct Vpr-mediated degradation are likely to provide an *in vivo* replicative advantage for all primate lentiviruses.

While none of the identified HIV-1 Vpr targets were degraded by the HIV-2 Vpx (HIV-2_ROD_) tested, we noted a shared ability of HIV-2 Vpx and SIVagm Vpr to deplete TASOR, a critical component of the Human Silencing Hub (HuSH) transcription repressor complex (Tchasovnikarova et al., 2015) (**Figure S5C**). Whilst this manuscript was in preparation, two other groups independently discovered and reported Vpx-mediated depletion of TASOR (Chougui et al., 2018; Yurkovetskiy et al., 2018). The HuSH complex mediates position-dependent transcriptional repression of a subset of lentiviral integrations, and we previously showed that antagonism of HuSH is able to potentiate HIV reactivation in the J-LAT model of latency (Tchasovnikarova et al., 2015). As predicted, Vpx-VLPs phenocopied the effect of RNAi-mediated TASOR depletion on reactivation of the HuSH-sensitive J-LAT clone A1 (**Figure S5D**).

Previous reports (Chougui et al., 2018; Yurkovetskiy et al., 2018) were conflicted on the conservation of TASOR depletion across Vpr variants. With some exceptions, most primate lentiviruses can be categorised into 5 lineages (**Figure S5E)**, of which two encode Vpx. The previously described canonical function of Vpx is the degradation of SAMHD1 (Hrecka et al., 2011; Laguette et al., 2011). Two lentiviral lineages lack Vpx, but use Vpr to degrade SAMHD1, while the HIV-1/SIVcpz lineage lacks SAMHD1 antagonism (Lim et al., 2012). We considered that TASOR antagonism may follow the same pattern, and thus tested Vpx proteins from both Vpx bearing lineages, and representative Vpr variants from lineages that use Vpr to degrade SAMHD1. All of these proteins were able to deplete TASOR in Vpx/Vpr transduced cells (**Figure S5F**). Antagonism of SAMHD1 and TASOR therefore follow the same pattern.

While the targeting of some substrates was conserved across Vpr varients from multiple lineages, we did not observe broad proteome remodelling outside the HIV-1/SIVcpz lineage. However, in the experiment described in **Figure 5A** an HIV-1 based lentiviral transduction system was used. Of the Vpr and Vpx variants tested, only Vpr proteins from the HIV-1/SIVcpz alleles are efficiently packaged. In these cases, cells receive both incoming and *de novo* synthesized Vpr, while in the case of other variants tested, only *de novo* synthesized Vpr is present. There is therefore a time-lag of at 18-24 hours for viral entry, reverse transcription, integration and de novo synthesis of protein to begin. As such, the more limited proteome remodelling seen in Vpr variants could reflect a lack of time for such changes to occur.

In order to account for this, we carried out an experiment in which cells were transduced with Vpr or Vpx from the primary HIV-2 isolate 7312a and assayed 48 hours to 96 hours post-transduction (**Figure 5E**), allowing time for additional changes to develop after *de novo* Vpr/Vpx synthesis. Even at 96 hours post transduction, HIV-2 Vpr showed very limited changes. The majority of these changes consisted of the depletion of proteins also targeted by HIV-1 Vpr (**Figure 5F** left panel).

Curiously, 7312a HIV-2 Vpx also depleted several proteins modulated by HIV-1 Vpr (**Figure 5F** – right panel), including direct targets for HIV-1 Vpr mediated degradation, BBX, HASPIN and ARHGAP11A. Some proteins were degraded by both HIV-2 Vpr and Vpx, while others were only degraded by the Vpx of this isolate. In lentiviral strains and/or lineages that encode both Vpr and Vpx, responsibility for degrading certain targets of HIV-1 Vpr is therefore shared between Vpr and Vpx, further emphasising their *in vivo* importance. Conversely, global proteome remodelling is therefore unique to Vpr variants of the HIV-1/SIVcpz lineage. This activity may help explain why viruses from this lineage appear to be more pathogenic than other primate lentiviruses (Greenwood et al., 2015; Keele et al., 2009), particularly given the potential to drive these changes in Vpr-exposed but uninfected bystander cells.

## Discussion

Proteomic analyses of cells infected with viruses from different orders have revealed widespread and varied changes to the cellular proteome (Diamond et al., 2010; Ersing et al., 2017; Greenwood et al., 2016; Weekes et al., 2014). While these changes are presumed to be multifactorial, in the case of HIV-1 infection, our data show that the majority of changes can be attributed to the action of a single viral protein, Vpr. We propose that this massive cellular proteome remodelling consist of firstly, the direct targeting of multiple cellular proteins for proteosomal degradation via the DCAF1/DDB1/CUL4A E3 ligase complex, followed by resulting secondary effects on other proteins. While many of the changes caused by Vpr are secondary, they occur within the physiological timeframe of productive infection (Murray et al., 2011; Perelson et al., 1996), and are therefore relevant to our understanding of the HIV-1-infected cell. Further, we have recently mapped changes to the cellular proteome in HIV-1 infected primary T-cells and confirmed that the Vpr-mediated changes in our CEM-T4 model represent the majority of changes in primary cells (Naamati et al., 2018).

Before this study, the list of direct Vpr targets was already extensive. We have confirmed here Vpr-mediated depletion of the previously described Vpr targets HLTF (Hrecka et al., 2016; Lahouassa et al., 2016), ZGPAT (Maudet et al., 2013), MCM10 (Romani et al., 2015), UNG2 (Schrofelbauer et al., 2005), TET2 (Lv et al., 2018), MUS81 and EME1 (Laguette et al., 2014; Zhou et al., 2016), while SMUG1 (Schrofelbauer et al., 2005) and PHF13 (Hofmann et al., 2017) were not detected in our model T-cell line. By analogy with other HIV accessory proteins, it might have been predicted that this list of Vpr targets would be nearly complete. Instead, we show here that it is only the tip of the iceberg.

While surprising, the ability of Vpr to degrade multiple cellular factors may be explained by the biology of this small protein. Mechanistically, depletion of multiple proteins with nucleic acid binding properties is consistent with known structural determinants of Vpr substrate recruitment. From the functional viewpoint, while HIV-1 has three accessory proteins (Vpu, Nef and Vif) to aid viral replication and counteract host defences in the late stages of the viral replication cycle, Vpr is the only HIV-1 accessory protein specifically packaged in virions. Multiple targets may therefore be required both to protect incoming virions from cellular factors, and to prime newly infected cells for productive viral replication.

This work does not represent the first attempt to use unbiased proteomics analysis to characterise Vpr function. In general, previous studies have used single proteomic experiments or methods to identify candidate proteins that either interact with, or are depleted by, Vpr, with individual proteins followed up using targeted immunoreagents. For example, Vpr binding partners were identified by Jager *et al.* (Jager et al., 2011) and Hrecka *et al.* (Hrecka et al., 2016) using IP-MS. Amongst these proteins, Jager *et al.* did not determine whether any were depleted. Conversely, Hrecka *et al.* (Hrecka et al., 2016) focussed on the single target protein, HLTF, and found it to be depleted by Vpr. Similarly, Lahuassa *et al.* (Lahouassa et al., 2016) used a SILAC-based approach to quantify proteomic changes in cells exposed to viral particles bearing Vpr, identifying 8 proteins which were depleted by at least 20%. Of these, only one, HLTF was confirmed to be a direct Vpr target by targeted orthogonal approaches.

As expected, the lists of ‘candidate’ Vpr proteins, identified, but not pursued, in the above studies overlaps with this work, and in some instances we have confirmed these candidates to be direct targets for Vpr mediated degradation. For example, SMN1 was found to bind Vpr by Jager *et al*, and to be degraded by Vpr by Lahuassa *et al.* in indepenedent proteomics experiments. Similarly, ESCO2 was found to bind Vpr by Hrecka *et al,* but the depletion was not confirmed by immunoblot – most likely due to poor performance of the commercial antibody used. We have shown here that both of these proteins are direct targets for Vpr-mediated degradation.

By contrast to previous studies, rather than using a targeted approach to follow-up only a small number of potential Vpr targets, we have combined complementary proteomic analyses to describe (i) the global proteome remodelling caused by Vpr, and (ii) multiple direct substrates for Vpr-mediated depletion. The two established criteria for Vpr targets, (binding and destabilisation) are satisfied by numerous proteins identified here, including ESCO2, SMN1, BBX and KIF18A. These proteins are therefore not candidate Vpr targets, but *bone fide* Vpr targets, proven to the same standard of evidence as other, previously described substrates.

However, in our proposed model, Vpr binds and degrades multiple cellular proteins, with the total pool of Vpr shared over multiple targets. The identification of cellular targets by co-IP is thus technically problematic and more prone to false negatives compared to other proteins that establish interactions with a small number of binding partners. We therefore used an alternative method of identifying direct targets for Vpr mediated depletion – proteins that are post-translationally degraded within 6 hours of treatment with Vpr. This strategy was more successful at capturing the known effects on cellular Vpr targets than the MS-IP approach, with the degradation of HLTF and ZGPAT being demonstrated within 6 hours of Vpr exposure. We are confident that the other proteins identified in this fashion also represent direct targets for Vpr mediated degradation, as secondary effects are excluded by both intrinsic elements of the technique, and the short time frame allowed.

A comparison of this work with the recent publication by Lapek *et al.* (Lapek et al., 2017) is required, given similarities in approach. Lapek *et al.* used an inducible HIV-1 provirus and quantify changes to the cellular proteome up to 24 hours post activation. They compared a wild type and a Vpr-deficient provirus, but find many fewer differences than in this study – indeed, they identified only nine proteins with significant difference between wild type and ΔVpr, none of which are previously defined Vpr targets – although DCAF1 is significantly reduced in the presence of Vpr. The difference most likely relates to differences between the systems used. In our model system, as in physiological conditions, Vpr is delivered with the virus particle at ‘0 h’ but de novo production of Vpr occurs late in the virus lifecycle. In Lapek *et al*, the experiment is concluded 24 hours after induction of the provirus. As production of de novo Vpr is Rev-dependent, Vpr production must occur after a significant delay within that 24-hour period. Thus, while some limited primary effects are evident, such as the depletion of DCAF1, other primary and secondary effects may not have had time to occur. Aside from the matter of timing, it is also worth noting that that in the Lapek *et al.* system, all Vpr is produced concurrently with Gag. The interaction between Vpr and Gag, and the recruitment of a proportion of Vpr into nascent viral particles, may inhibit nuclear localisation and substrate degradation.

Despite the importance of Vpr *in vivo*, an *in vitro* viral replication phenotype is often absent in T-cell infection models (Guenzel et al., 2014). Nonetheless, expression of Vpr in T-cells causes cell cycle arrest (Bolton and Lenardo, 2007; Gummuluru and Emerman, 1999; Rogel et al., 1995), cell death (Bolton and Lenardo, 2007), transactivation of the viral LTR (Gummuluru and Emerman, 1999), enhancement or antagonism of crucial signalling pathways including NFAT (Hohne et al., 2016) and NF-κB (Liang et al., 2015; Liu et al., 2014; Muthumani et al., 2006), disruption of PARP1 localisation (Hohne et al., 2016; Muthumani et al., 2006), defects in chromatid cohesion (Shimura et al., 2011) and induction of the DNA damage response (Richard et al., 2010; Vassena et al., 2013). At least some of these phenotypes can be segregated (Bolton and Lenardo, 2007; Hohne et al., 2016). The molecular mechanisms underpinning these phenomena have remained controversial. In our model, we propose that these multiple phenotypes can be explained by Vpr targeting multiple cellular proteins and pathways, with potential for redundant or cumulative effects.

Here, we have considered the most well described cellular phenotype for Vpr, G2/M cell cycle arrest, with findings compatible with this model. In addition to confirming the previously described effect of Vpr mediated MCM10 degradation, we have identified three other proteins that are directly targeted by Vpr, show depletion correlating with the extent of G2/M mediated arrest in a panel of Vpr mutants and result in arrest at G2/M when depleted through RNAi. Notably, depletion of two of these proteins, SMN1 and CDCA2 (Repo-Man), has been shown to activate the DNA damage response and stimulate ATM/ATR kinase activity (Kannan et al., 2018; Peng et al., 2010), demonstrated by many groups to be a critical step towards the G2/M arrest caused by Vpr (Berger et al., 2015; Fregoso and Emerman, 2016; Roshal et al., 2003). The contribution of multiple Vpr targets towards the same cellular phenotype may also be exemplified by another described cellular phenotype for Vpr, premature chromatid segregation (PCS) (Shimura et al., 2011). While not specifically investigated here, RNAi-mediated knockdown of three proteins depleted by Vpr, ESCO2, CDCA5 (Sororin) and HASPIN, have been previously associated with this phenotype in different systems (Dai et al., 2006; Hou and Zou, 2005; Rankin et al., 2005).

In conclusion, Vpr degrades multiple cellular targets, resulting in global remodelling of the host proteome, and labyrinthine changes to different cellular pathways. This explains why its effects on cellular phenotypes and viral replication are complex and remain poorly understood, why the functional consequences of individual Vpr targets identified and studied in isolation have proved elusive, and why the search for a single, critical Vpr target has been problematic.

## Acknowledgements

This work was supported by the Wellcome Trust (PRF 101835/Z/13/Z to PJL), the MRC (CSF MR/P008801/1 to NJM), NHSBT (WPA15-02 to NJM), the NIHR Cambridge BRC, and a Wellcome Trust Strategic Award to the CIMR. The authors thank Dr. Reiner Schulte and the CIMR Flow Cytometry Core Facility team, and the Lehner laboratory for critical discussion.

## Data availability

In addition to **Figure 1—source data 1**, which includes data from all of the proteomics experiments carried out here, proteomics data have been deposited to the ProteomeXchange Consortium via the PRIDE (Vizcaino et al., 2016) partner repository with the dataset identifier PXD01029.

## Experimental Procedures

### NL4-3 molecular clones

pNL4-3-dE-EGFP (derived from the HIV-1 molecular clone pNL4-3 but encoding Enhanced Green Fluorescent Protein (EGFP) in the *env* open reading frame (ORF), rendering Env non-functional) was obtained through the AIDS Reagent Program, Division of AIDS, NIAD, NIH: Drs Haili Zhang, Yan Zhou, and Robert Siliciano (Zhang et al., 2004) and the complete sequence verified by Sanger sequencing. The ΔVpr mutant was generated by cloning three stop codons into the Vpr open reading frame, immediately after the overlap with Vif to prevent interference with that gene:

WT Vpr ORF sequence

ATGGAACAAGCCCCAGAAGACCAAGGGCCACAGAGGGAGCCATACAATGAATGGACACTAGAGCTTTTAGAGGAA

ΔVpr ORF sequence

ATGGAACAAGCCCCAGAAGACCAAGGGCCACAGAGGGAGCCATACAATGAATGGACACTAGAGCTT<u>TAATAGTAA</u>

### Viral stocks

VSVg-pseudotyped NL4-3-dE-EGFP HIV viral stocks were generated by co-transfection with pMD.G (VSVg) as previously described (Greenwood et al., 2016). NL4-3-dE-EGFP HIV viral stocks were titred by infection/transduction of known numbers of relevant target cells under standard experimental conditions followed by flow cytometry for GFP and CD4 at 48 hr to identify % infected cells. Single Vpr and Vpx proteins were expressed in a modified dual promotor pHRSIN (van den Boomen et al., 2014) vector in which Vpr or Vpx expression is driven by the rous sarcoma virus (RSV) promotor, and Emerald GFP expression is driven by the ubiquitin promotor. Virus was generated by co-transfection with p8.91 and pMD.G in 293T as previously described (Greenwood et al., 2016). Infectious MOI was normalised by infection in the absence of reverse transcription (RTi) inhibitors, which were included where specified (see below). The panel of Vpr and Vpr proteins used are detailed in **Table 2**. Amino acid sequences were codon optimized and synthesized as double stranded DNA (IDT), inserted into the empty vector construct by Gibson assembly, and confirmed by sanger sequencing.

### CEM-T4 T-cell infections

CEM-T4 T-cells were infected with concentrated NL4-3-dE-EGFP or pHRSIN lentiviral stocks by spinoculation at 800 ×g for 1 h in a non-refrigerated benchtop centrifuge in complete media supplemented with 10 mM HEPES. Where reverse transcription treatment was specified (RTi), cells were incubated with zidovudine (10 μM) and efavirenz (100 nM) (AIDS Reagent Program, Division of AIDS, NIAD, NIH) for 1 hr prior to spinoculation, and inhibitors maintained at these concentrations during subsequent cell culture. For MS experiments, cells were subject to dead cell removal (magnetic dead cell removal kit, Miltenyi). Subsequent sample preparation, TMT labelling, and MS are described below, at the end of this section.

### Antibodies

Antibodies against the following proteins were used for immunoblot, listed by manufacturer: Atlas antibodies: TASOR (HPA006735). Bethyl: BBX (A303-151A), HLTF (A300-230A), RALY (A302-070A), ZNF512B (A303-234A). Cell signalling : SMN1/2 (2F1). Novus: ESCO2 (NB100-87021). Origine: UNG2 (2C12). Proteintech: Vpr (51143-I-AP). Santa Cruz: CCNB1 (SC-245), ZGPAT (SC-51524). Sigma: β-actin (AC74). Abcam : p24 (ab9071), VCP (ab11433). The following secondary antibodies were used: goat anti-mouse-HRP and anti-rabbit-HRP (immunoblot, Jackson ImmunoResearch, West Grove, PA). Anti-CD4-AF647 (clone OKT4; BioLegend) and anti-CD271 (NGFR)-APC (clone ME20.4, Biolegend) were used for flow cytometry.

### Statistical analysis

Anova and Fisher’s exact test analysis were carried out as described in figure legends using Graphpad Prism (v7.04). TMT multiplexed proteomics datasets were analysed using a moderated T-test analysis with Benjamini-Hochberg correction (Schwammle et al., 2013), carried out using R (v3.3.1) (R Core Team, 2013). Significance B values for pulsed-SILAC analysis were calculated using Perseus (Tyanova et al., 2016).

Further details can be found in the supplemental experimental procedures.

## Supplemental Information

**Table S1. Interactive database of protein changes in the datasets presented here**

**Figure S1 – Related to Figure 2.**
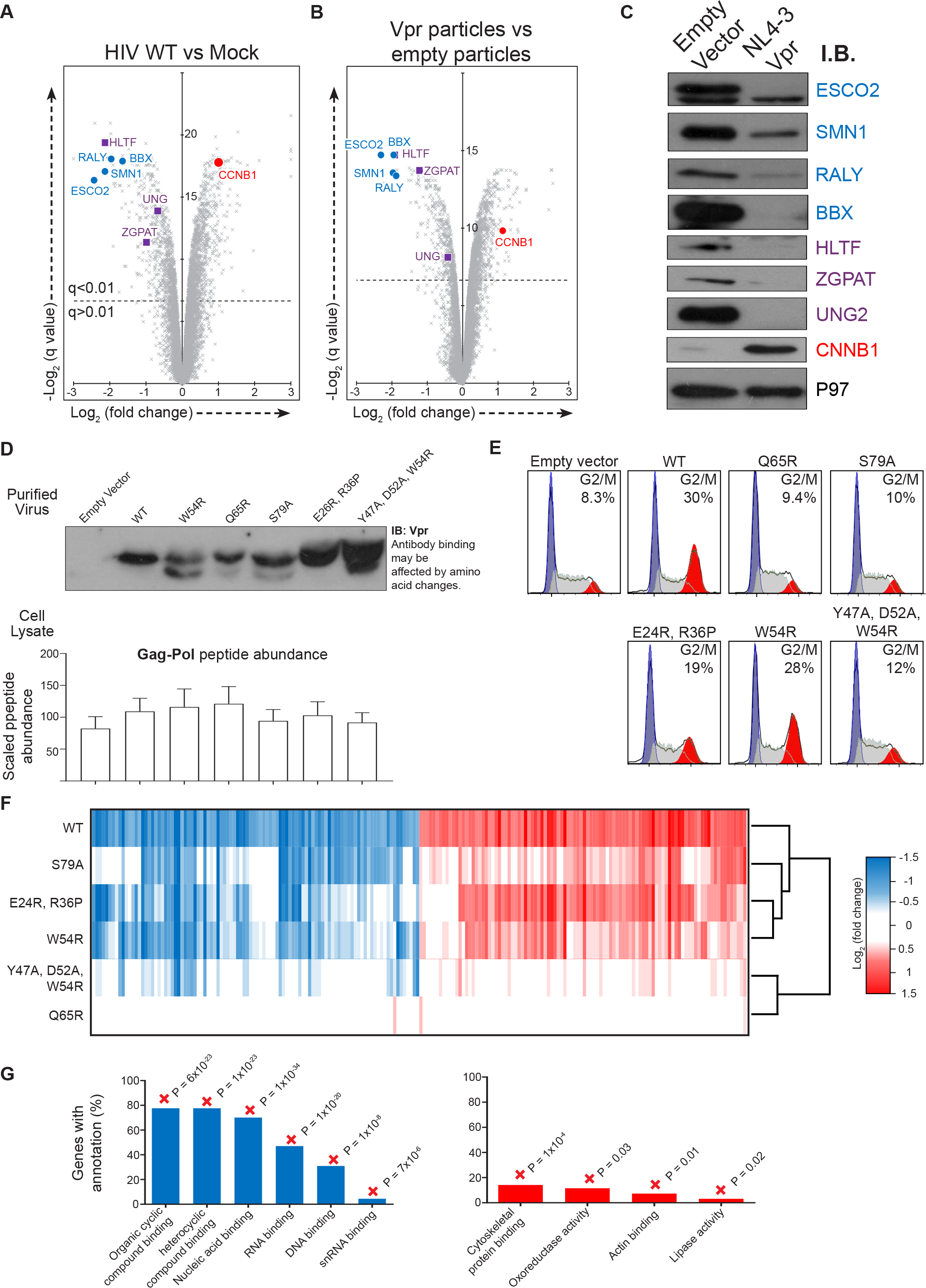
Immunoblot replication of proteins altered by Vpr and additional information relating to experiment described in Figure 2. **A,** Scatterplots displaying pairwise comparisons between WT, ΔVpr and mock-infected cells. Vpr depleted proteins confirmed by blot in this figure are highlighted in blue, UNG, ZGPAT and HLTF are previously identified targets for Vpr mediated degradation, shown in purple. CCNB1 is increased by Vpr (red). **B,** Scatterplot displaying pairwise comparison between cells exposed to empty or Vpr bearing viral particles. **C,** Immunoblot of proteins highlighted in this figure. **D,** Immunoblot of purified virus preparations used to infect cells for the proteomics experiment displayed in **Fig 2A**. **B,** Bar chart showing the average scaled abundance of matrix, capsid and integrase peptides detected in the cell lysate by MS. Bars show mean and standard deviation. **E,** 7-AAD stain of cells exposed to empty vector, or Vpr wild type or Vpr mutants. Watson pragmatic modelling was used to identify cells in G1 (blue), S (grey) or G2/M (red) phase. **F**, Heatmap showing the behaviour of the 100 proteins most depleted by Vpr particles (blue) and increased (red) within the defined highly modulated subsets. Colour indicates the log_2_ fold change of each protein in each condition compared to empty particle treatment. Genes were clustered using uncentered Pearson correlation and centroid linkage, and conditions clustered by column means. **G,** Defined groups of Vpr depleted and increased proteins were subject to gene ontology enrichment analysis, compared with a background of all proteins quantitated in these experiments. Plots show all significantly enriched (Fisher’s exact test with Bonferroni correction) molecular function terms with associated p values. + / − indicates if the classification was enriched or de-enriched compared with the expected number of proteins expected by chance. Where significant, associated p value represents the results of a Fisher’s exact test with Bonferroni correction. This is highly conservative as it is corrected for all 1061 possible cellular compartment terms, not just those shown.

**Figure S2 – Related to Figure 2.**
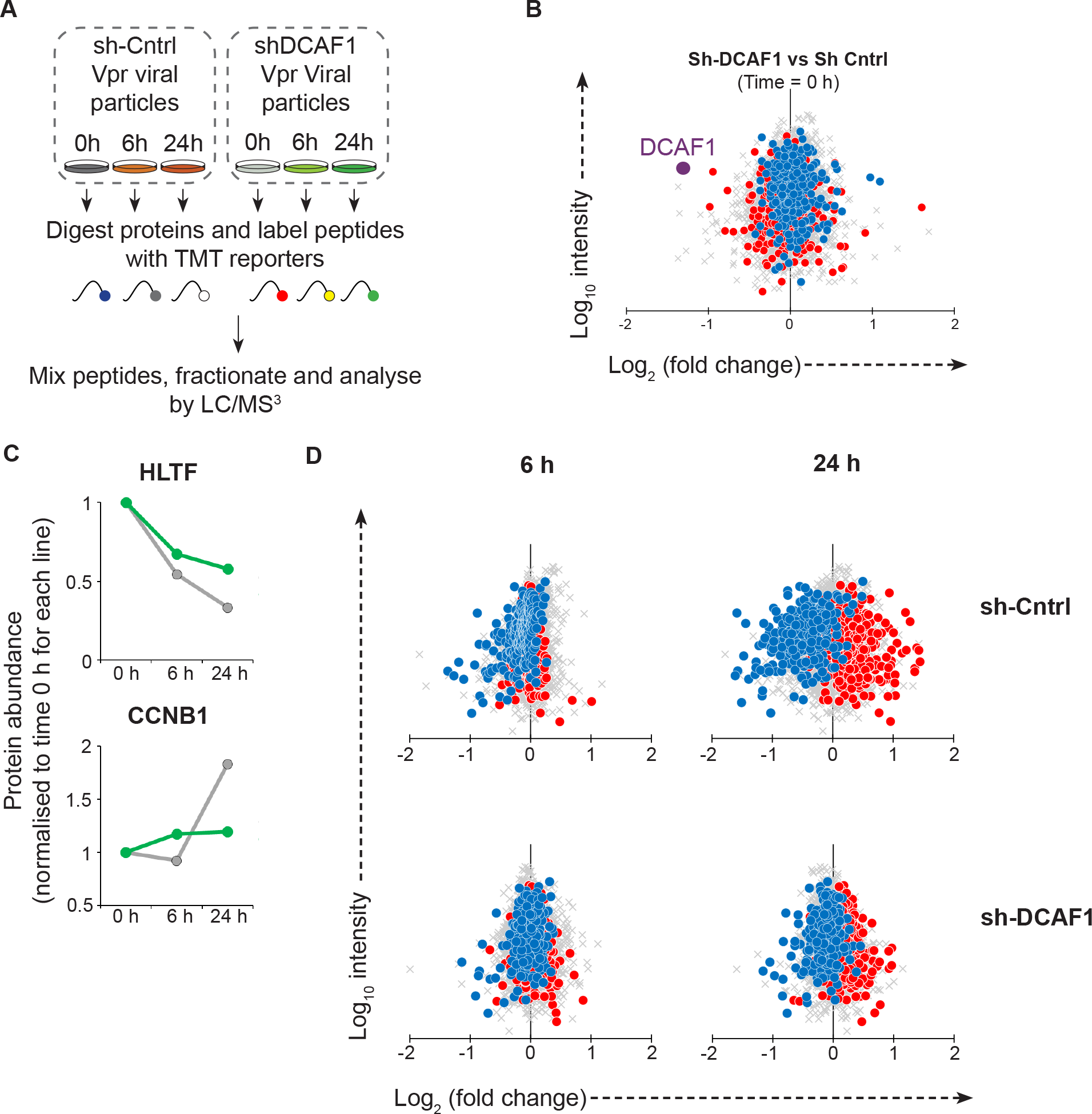
Quantifying the effect of Vpr under reduced DCAF1 conditions. **A,** Graphical summary of the DCAF1 KD experiment. **B,** Scatterplot displaying pairwise comparison between Sh-Control and Sh-DCAF1 cells at 0 h, with defined groups of 302 Vpr depleted (blue) and 413 increased (red) proteins highlighted. DCAF1 is highlighted in purple. **C,** Example time-course behaviour for one Vpr target (HLTF) and one secondary Vpr effect (CCNB1). **D,** Scatterplots showing the pairwise comparison between the 6 or 24 h time-point with the 0 h condition for each sh-transduced cell line.

**Figure S3 – Related to Figure 2.**
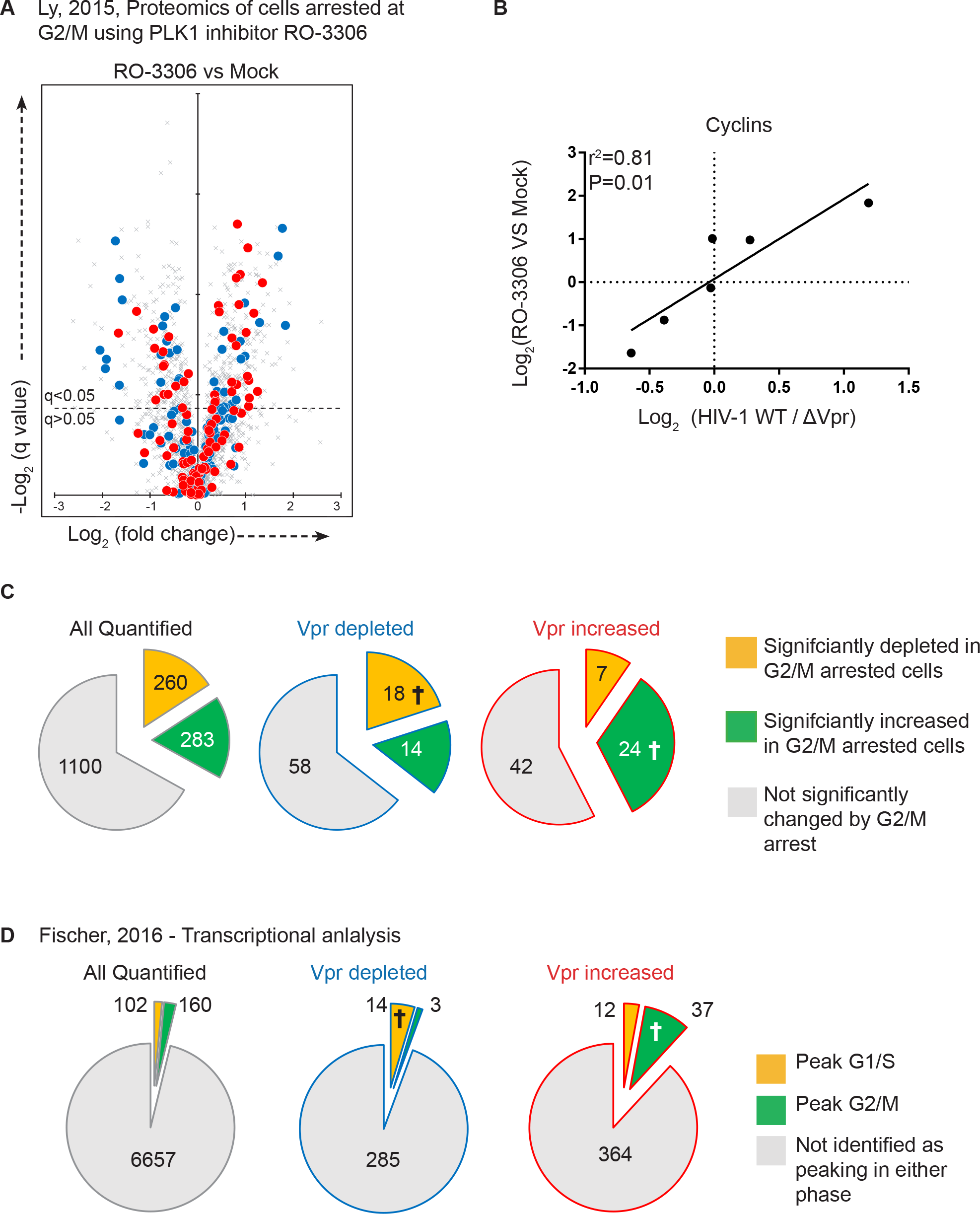
Cell cycle regulation of Vpr modulated proteins. **A,** Ly, 2015 contains a proteomic analysis of NB4 cells treated with RO-3306 cells arrested in G2/M with the PLK1 inhibtor RO-3306. Scatterplot showing the pairwise comparison of abundance of proteins isolated from RO-3306 treated vs mock treated cells. Groups of proteins defined in the current study of 302 Vpr depleted (blue) and 413 increased (red) proteins are highlighted, indicating the behaviour of these proteins in NB4 cells arrested at G2/M. **B,** Correlation of the Vpr mediated change in cyclin abundance in the present study (x-axis), with RO-3306 mediated changes in NB4 cells in Ly, 2015. Changes in cyclin abundance in HIV infection are assumed to be secondary to cell cycle arrest, and thus this correlation indicates the concordance between effects secondary to cell cycle arrest between the two datasets. **C,** Pie charts showing the overlap between changes in the present study and changes induced by G2/M arrest in Ly, 2015. Left panel shows the behaviour of all proteins quantified in both the present study (**Figure 1A** and **Figure 2A**) and Ly, 2015, i.e. a total of 1643 proteins were quantified in both datasets, of which 1100 proteins did not significantly change in G2/M arrest, 260 proteins were significantly depleted in G2/M arrested cells, and 283 proteins were significantly increased in G2/M arrested cells. Middle panel shows the behaviour the defined group of 302 Vpr depleted proteins. I.e. of 302 proteins, a total of 90 proteins were detected in Ly, 2015, 58 of which did not significantly change in G2/M arrest, 18 were significantly depleted in G2/M arrest and 14 were significantly increased in G2/M arrest. † Indicates the fraction in each case where the change induced by cell cycle arrest is in the same direction as the effect of Vpr, i.e. the fraction for which cell cycle arrest could explain the Vpr mediated protein changes. Right panel shows the behaviour of the defined group of 413 Vpr increased proteins. **D,** Fischer, 2016, define lists of 115 proteins whose expression peaks in G1/S phase and 174 proteins whose expression peaks in G2/M phase. Pie-charts show the overlap between these lists and (left) all proteins detected in the present study, (middle) proteins defined as being depleted by Vpr, and (right) proteins defined as being increased by Vpr. As in the proteomics dataset, there is some enrichment of proteins with peak expression in G2/M within Vpr increased proteins, and some enrichment of proteins with peak expression in G1/S in Vpr depleted proteins, consistent with some effects being secondary to cell cycle arrest, but these effects are in the minority.

**Figure S4 – Related to Figure 2.**
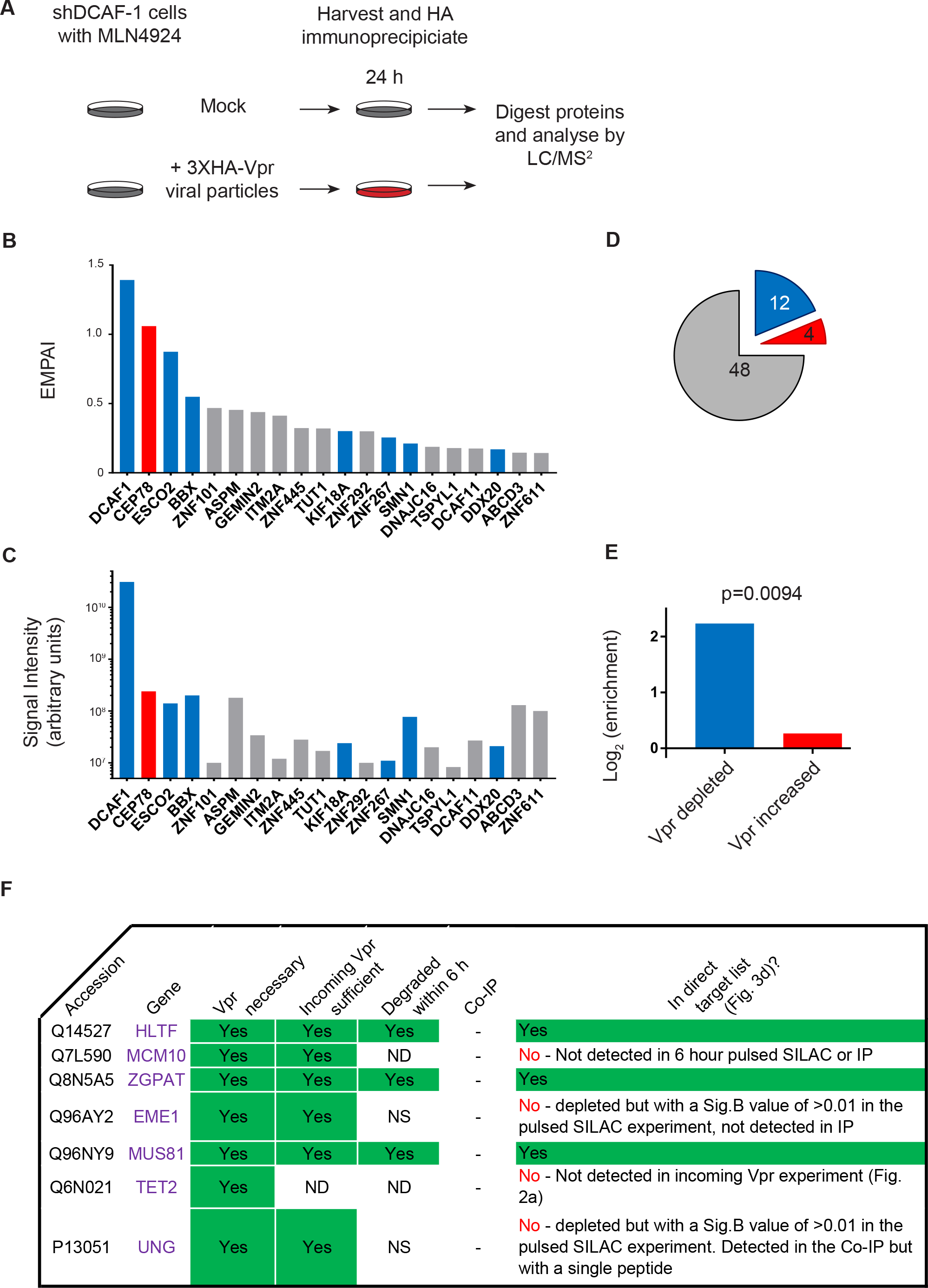
MS-IP approach to identify direct targets for Vpr mediated degradation. **A,** Graphical summary of the MS-IP experiment. All cells were stably transduced with an ShDCAF1 vector as described earlier. MLN4924 is a pan-Cullin inhibitor. **B,** 20 most abundant proteins identified by Co-IP determined by number of unique peptides, normalised as a proportion of the maximum possible peptide count for each protein, (Exponentially modified protein abundance index, emPAI). Proteins falling within the defined list of 302 Vpr depleted and 413 Vpr increased proteins are highlighted in blue and red respectively. **C,** The same 20 proteins with signal intensity rather than peptide number shown. **D,** Pie chart indicating the overlap between the proteins co-immunoprecipitated with Vpr and the defined list of 302 Vpr decreased (blue) and 413 Vpr increased proteins (red), and proteins detected but falling into neither list (grey). **E**, Bar chart showing the enrichment of Vpr depleted and Vpr increased proteins within proteins co-immunoprecipitated with Vpr compared to the expected numbers of proteins that would be co-immunopreciptated from each group by chance. p value indicates a Fisher’s exact test of a 2×2 contingency table (Vpr depleted/increased, identified by co-IP/not identified), indicating that the two criteria are significantly linked.

**Figure S4 – Related to Figure 5.**
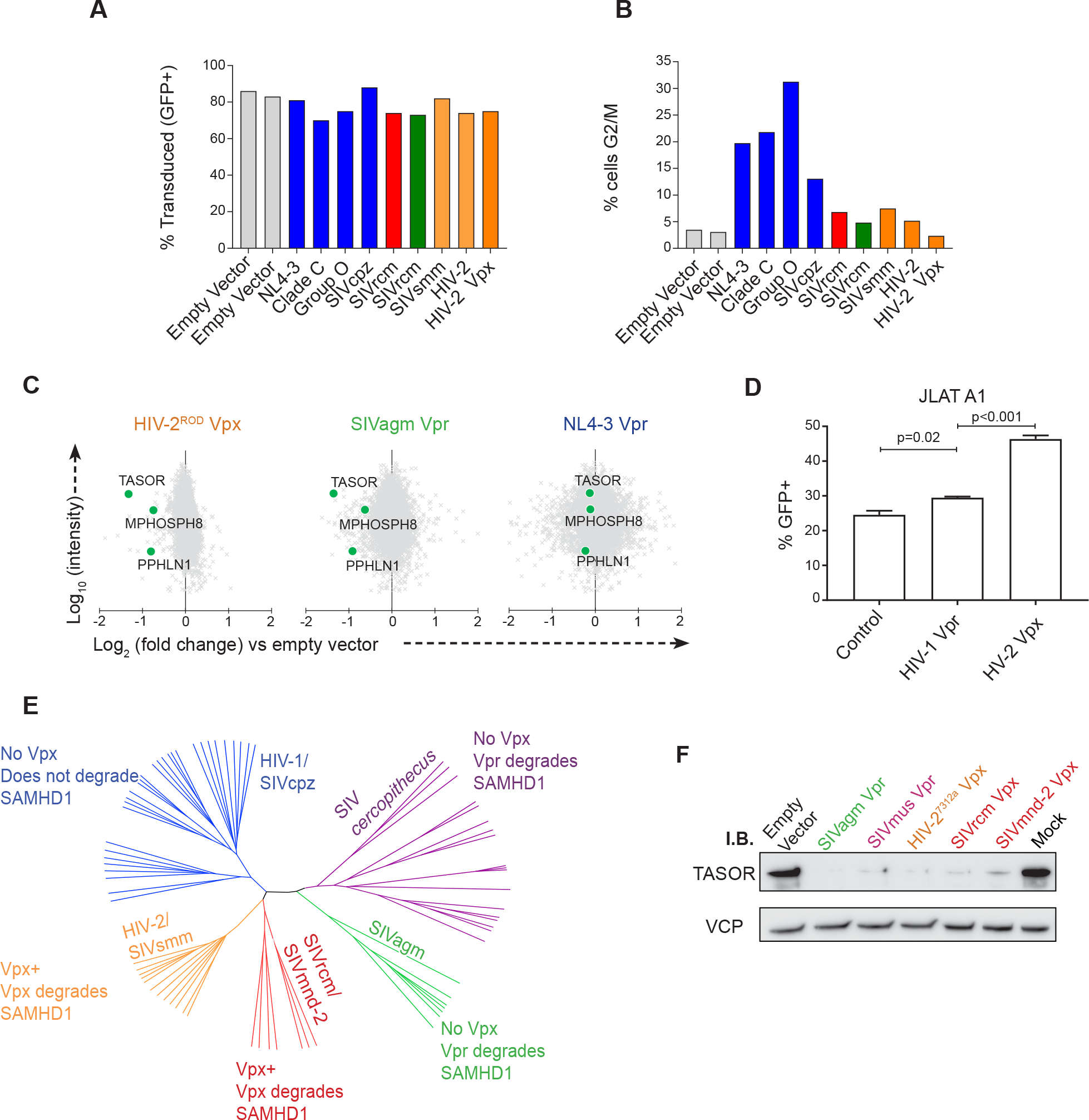
Additional information relating to expanded panel of primate lentiviral Vpr and Vpx varients & Depletion of HuSH components by lentiviral Vpx and Vpr proteins. **A,** %GFP positive (transduced) cells at harvest. Cells were transduced at an infectious MOI of 1.5 based on prior titration, with the actual resulting % transduction varying slightly across the samples. **B**, Proportion of cells in G2/M at point of harvest based on 7-AAD staining and Watson pragmatic modelling. **C,** Scatterplots showing the pairwise comparison between each Vpr tested and empty vector control with HuSH complex components highlighted. **D**, Bar graph of GFP percentage positive of JLAT-A1 cells after transduction with control (Cre recombinase), Vpr or Vpx proteins and treatment with TNFα. Mean and SEM of 3 biological replicates per condition are shown, representative of three independent similar experiments. p-values determined by ordinary one-way ANOVA with Bonferroni comparison between Vpr/Vpx treatment and control treated cells. **E,** Phylogenetic tree of primate lentiviruses based on an alignment of Vif nucleic acid sequences, with 5 major lineages of primate lentiviruses labelled. Information on Vpr and Vpx activity based on a selected number of isolates tested in each lineage in Lim, 2012 (Lim et al., 2012) **F,** Immunoblot of TASOR in cells transduced with a panel of Vpx and Vpr proteins.

**Table S2 – Related to Figure 5.**
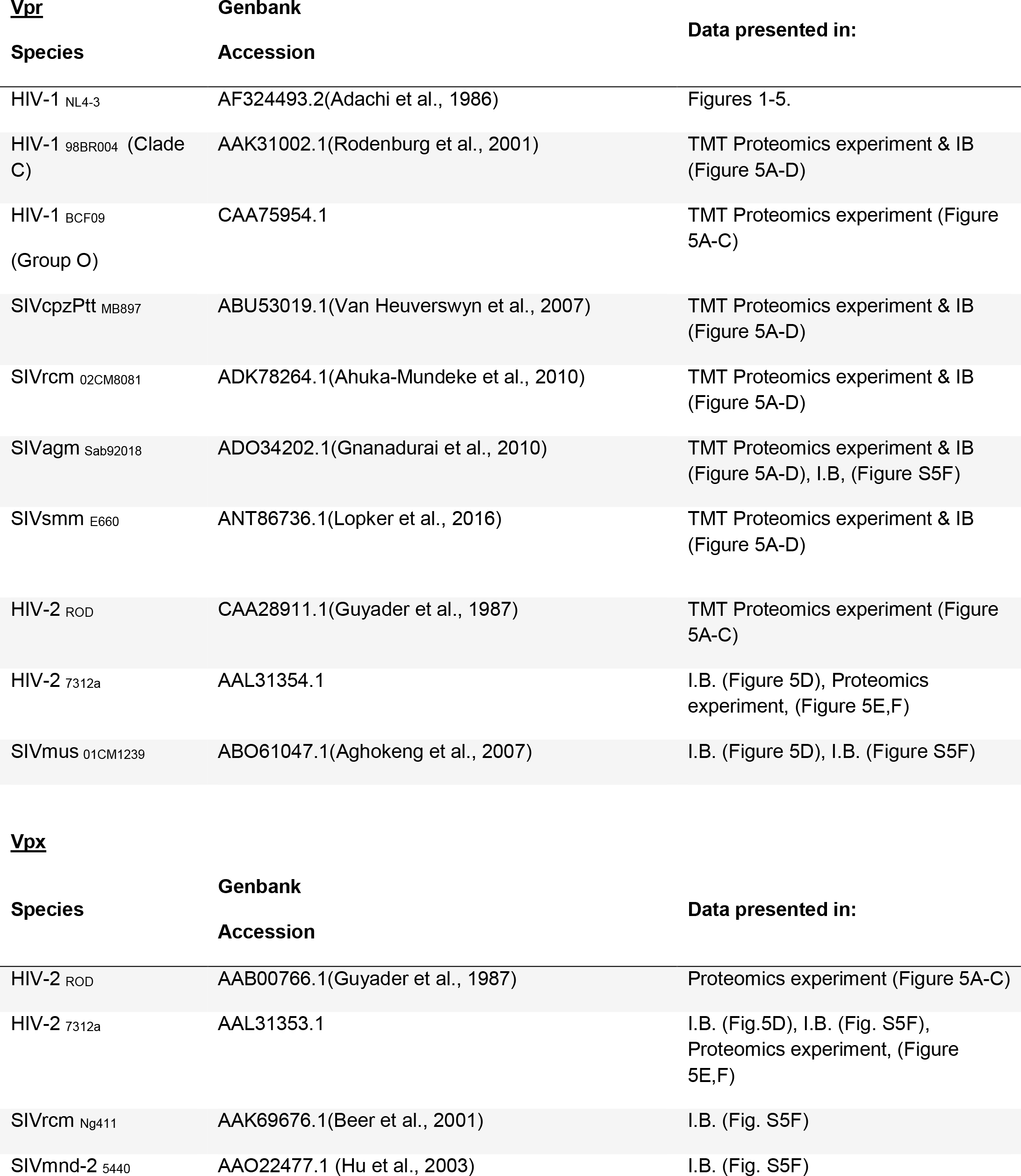
Vpr and Vpx variants used in this study.

| **<u>Vpr</u>** | **Genbank** | **Data presented in:** |
| --- | --- | --- |
| **Species** | **Accession** |  |
| HIV-1<sub>NL4-3</sub> | AF324493.2(Adachi et al., 1986) | Figures 1-5. |
| HIV-1<sub>98BR004</sub> (Clade C) | AAK31002.1(Rodenburg et al., 2001) | TMT Proteomics experiment & IB (Figure 5A-D) |
| HIV-1<sub>BCF09</sub><br>(Group O) | CAA75954.1 | TMT Proteomics experiment (Figure 5A-C) |
| SIVcpzPtt<sub>MB897</sub> | ABU53019.1(Van Heuverswyn et al., 2007) | TMT Proteomics experiment & IB (Figure 5A-D) |
| SIVrcm<sub>02CM8081</sub> | ADK78264.1(Ahuka-Mundeke et al., 2010) | TMT Proteomics experiment & IB (Figure 5A-D) |
| SIVagm<sub>Sab92018</sub> | ADO34202.1(Gnanadurai et al., 2010) | TMT Proteomics experiment & IB (Figure 5A-D), I.B. (Figure S5F) |
| SIVsmm<sub>E660</sub> | ANT86736.1(Lopker et al., 2016) | TMT Proteomics experiment & IB (Figure 5A-D) |
| HIV-2<sub>ROD</sub> | CAA28911.1(Guyader et al., 1987) | TMT Proteomics experiment (Figure 5A-C) |
| HIV-2<sub>7312a</sub> | AAL31354.1 | I.B. (Figure 5D), Proteomics experiment, (Figure 5E,F) |
| SIVmus<sub>01CM1239</sub> | ABO61047.1(Aghokeng et al., 2007) | I.B. (Figure 5D), I.B. (Figure S5F) |

**Vpx**
| **Species** | **Genbank**<br>**Accession** | **Data presented in:** |
| --- | --- | --- |
| HIV-2<sub>ROD</sub> | AAB00766.1(Guyader et al., 1987) | Proteomics experiment (Figure 5A-C) |
| HIV-2<sub>7312a</sub> | AAL31353.1 | I.B. (Fig.5D), I.B. (Fig. S5F),<br>Proteomics experiment, (Figure 5E,F) |
| SIVrcm<sub>Ng411</sub> | AAK69676.1(Beer et al., 2001) | I.B. (Fig. S5F) |
| SIVmnd-2<sub>5440</sub> | AAO22477.1 (Hu et al., 2003) | I.B. (Fig. S5F) |

### Supplemental Experimental Procedures

#### Lentivector for shRNA expression

For lentiviral shRNA-mediated knockdown of DCAF1 hairpins were cloned into pHRSIREN-PGK-hygro, with transduced cells selected for hygromycin resistance.

The following oligonucleotide was inserted using BamHI-EcoRI (only top oligonucleotide shown), identified from the Broad Institute GPP Web portal. Gene specific target sequence (forward and reverse complement) is underlined.

#### DCAF1

GATCC<u>GCTGAGAATACTCTTCAAGAA</u>TTCAAGAGA<u>TTCTTGAAGAGTATTCTCAGC</u>TTTTTTG

The same methodology was used to clone other shRNA haripins, the forward orientation targeting sequence used for each is shown below:

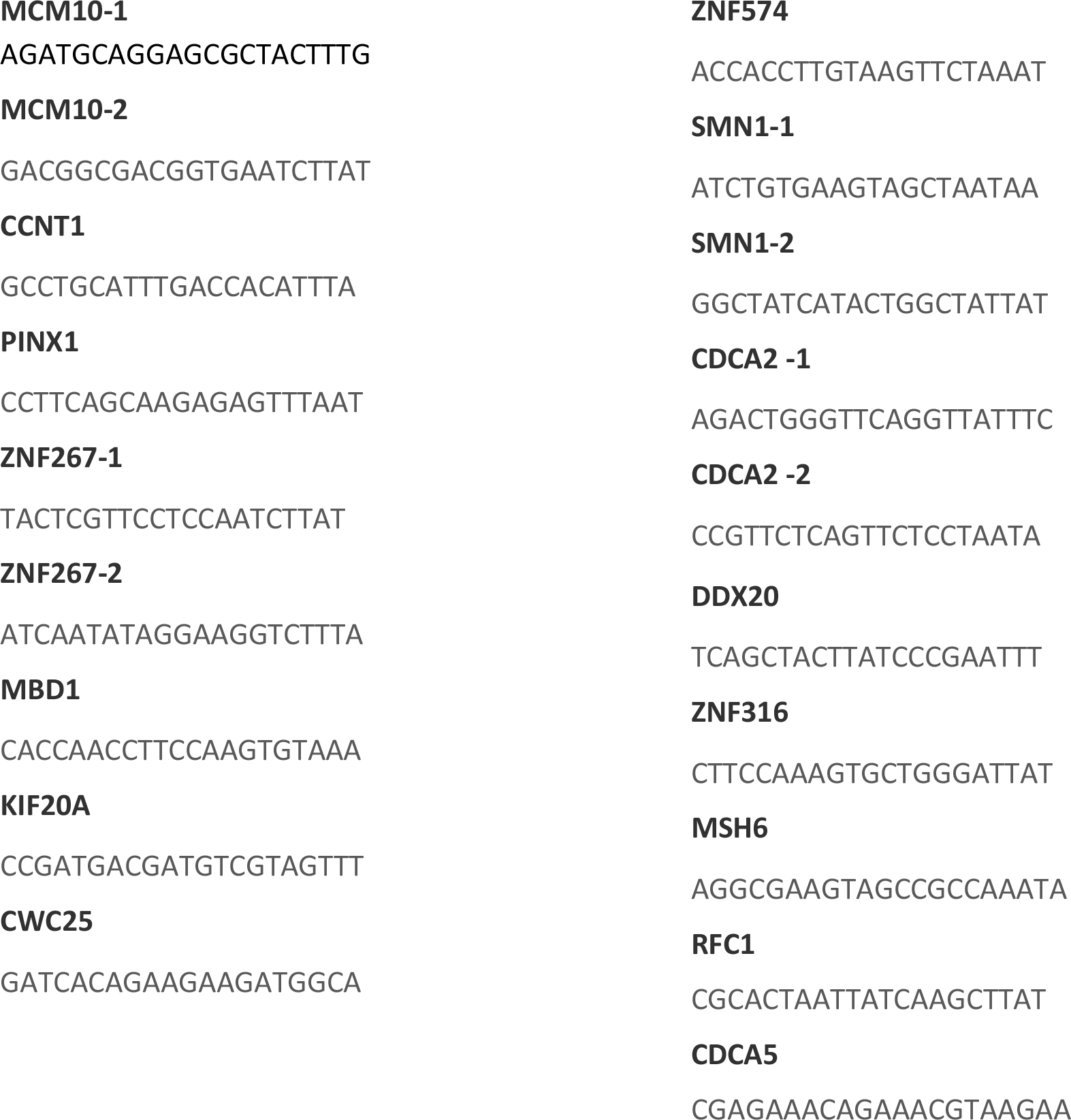

#### Gene ontology enrichment analysis

Analysis was carried out using Panther (Mi et al., 2017), using the web interface at: http://pantherdb.org/. Lists of proteins highly significantly depleted or increased by Vpr were compared to a background list of proteins quantified in the two experiments used to define those lists.

#### 7-AAD staining

Cells were fixed in ice cold 70% ethanol for at least 30 minutes, washed with PBS and stained in 25ug/ml 7-AAD for 15 minutes before acquisition on a BD FACSCalibur. Analysis was carried out in Flowjo, v.10.5.2. Dead cells and doublets were excluded by gating on forward scatter, side scatter, and fluorescent area and height. Univariate cell cycle modelling was carried out using the Watson pragmatic method.

#### ShRNA knockdown confirmation by RT-PCR

Total RNA was extracted using the RNeasy Plus Mini kit (Qiagen). Total RNA (250 ng) was reverse transcribed into cDNA using a poly(d)T primers and SuperScript III Reverse Transcriptase (Invitrogen) following the manufacturer's instructions. Real-time qRT–PCR was performed using the ABI 7500 Real-Time PCR system (Applied Biosystems) and SYBR Green PCR Master Mix (Applied Biosystems), with cycling parameters of 50°C for 2 min and 95°C for 5 min, followed by 40 cycles of 95°C for 15 s and 60°C for 1 min. The gene-of-interest specific primer pairs were predesigned and purchased from Sigma-Aldrich as KiCqStart primers, with the following sequences: MCM10_FOR 5′-CTTATACAGAAGAGGCTGATG and MCM10_REV 5’-CCTCTTGCAACTCTTCATTC; ZNF267_FOR 5’-GTAGAATTCTCTTTGGAGGAG and ZNF267_REV 5’-CTCACTCTTCACATTCCAAG; CDCA2_FOR 5’-AGGAAAGTCATCATCCTACC and CDCA2_REV 5’-GATGGTTTGTTTCAGGAGAG; SMN1_FOR 5’-GGAAAGCCAGGTCTAAAATTC and SMN1_REV 5’-AGAATCTGGACATATGGGAG. The difference in the amount of input cDNA was normalized to an internal control of GAPDH, using the following primers: GAPDH_FOR: 5’ ATGGGGAAGGTGAAGGTCG and GAPDH_REV: 5′-CTCCACGACGTACTCAGCG.

#### Mass spectrometry Immunoprecipitation (MS-IP)

CEM-T4 T-cells were lysed in 1 % NP-40. Lysates were pre-cleared with IgG-Sepharose (GE Healthcare, UK) and incubated for 3 hr at 4°C with anti-HA coupled to agarose beads (Sigma EZview Red Anti-HA Affinity Gel). After washing in 0.5% NP-40, samples were eluted with 0.5 mg/ml HA peptide at 37°C for 1 hr. Additional detail on MS methods is available at the end of this section. Proteins defined as co-immunoprecipitating with Vpr were detected with at least 3 peptides in the Vpr condition, not identified in the control condition, and were present in <20% of MS-IP available in the Crapome (Mellacheruvu et al., 2013) database http://crapome.org/.

#### Pulsed Stable isotope labelling with amino acids in cell culture (Pulsed-SILAC)

For SILAC labelling, CEM-T4 T-cells were grown for at least 7 cell divisions in SILAC RPMI lacking lysine and arginine (Thermo Scientific, Thermo Fisher Scientific, UK) supplemented with 10% dialysed FCS (Gibco, Thermo Fisher Scientific), 100 units/ml penicillin and 0.1 mg/ml streptomycin, 280 mg/L proline (Sigma, UK) and medium (K4, R6; Cambridge Isotope Laboratories, Tewksbury, MA) or heavy (K8, R10; Cambridge Isotope Laboratories) ^13^C/^15^N-containing lysine (K) and arginine (R) at 50 mg/L. At 0 h, cells were washed in media containing only light (^12^C /^14^N) lysine and arginine and were maintained in this media for the duration of the experiment. Additional detail on MS methods is available at the end of this section.

#### J-LAT reactivation experiments

JLAT A1 cells were transduced with Cre recombinase, NL4-3 Vpr or HIV-2 ROD Vpx within a pHRSIN IRES NGFR vector (Matheson et al., 2014). 24 h after transduction, cells were treated with 2 ng/ml TNFα (PeproTech, 300-01A). After an additional 24 h cells were stained with anti-NGFR APC and analysed for APC and GFP expression by flow cytometry. Cells were gated for NGFR+ cells to exclude non-transduced cells. Flow cytometry data was acquired on a BD FACSCalibur.

#### Phylogenetic tree of Vpr sequences

An existing nucleic acid sequence alignment of 191 representative Vif sequences from HIV-1, HIV-2 and SIVs from 22 primate species was downloaded from: https://www.hiv.lanl.gov/content/sequence/NEWALIGN/align.html

A tree was generated using the percentage identity average distance in jalview (Waterhouse et al., 2009) and visualised in Figtree: http://tree.bio.ed.ac.uk/software/figtree/

10 sequences from 5 primate species not falling into the 5 highlighted linages are not shown.

#### Sample Preparation for Mass spectrometry

Samples were prepared using three different methods depending on the experiment. Initial infection experiment was by SDC-FASP. VLP and shRNA experiments were by PreOmics NHS-iST sample preparation Kit. pSILAC and MS-IP experiments were by SP3.

##### SDC-FASP

Samples prepared essentially according to the protocol in Leon *et al*(Leon et al., 2013). Briefly samples were lysed in 50mM TEAB (pH8.5) 2% SDS, reduced and alkylated with TCEP/Iodoacetamide, quantified by BCA assay. 50ug of each sample was diluted with TEAB/8M urea for loading onto 30kDa ultrafiltration devices. Samples were washed 3 times with 500uL urea buffer and 3 times with digestion buffer (TEAB/0.5% sodium deoxycholate) before resuspending in 50uL digestion buffer containing 1ug trypsin and incubating overnight at 37 degrees. After digestion samples were spun through the filters and filters washed with 50uL TEAB. SDC was removed by acidification and two phase partitioning with ethyl acetate before vacuum drying and labelling with TMT reagents according to the manufacturer’s instructions.

##### PreOmics iST

Samples lysed in kit lysis buffer and quantified by BCA assay. 25ug of each sample was digested essentially according to manufacturer’s instructions, scaling volumes for a digestion of 25ug total protein.

##### SP3

Samples were lysed in 50mM TEAB (pH8.5) 2% SDS (MS-IPs were adjusted to 2% SDS), reduced and alkylated with TCEP/Iodoacetamide before digestion using the SP3 method(Hughes et al., 2014). Briefly, carboxylate modified paramagnetic beads are added to the sample and protein is bound to the beads by acidification with formic acid and addition of acetonitrile (ACN, final 50%). The beads are then washed sequentially with 100% ACN, 70% Ethanol (twice) and 100% ACN. 10-20uL TEAB (Triethylammonium bicarbonate) pH8 and 0.1% Sodium deoxycholate (SDC) is then added to the washed beads along with trypsin. Samples were then incubated overnight at 37 degrees with periodic shaking at 2000rpm. After digestion, peptides are immobilised on beads by addition of 200-400uL ACN and washed twice with 100uL ACN before eluting in 19uL 2% DMSO and removing the eluted peptide from the beads.

#### Off-line high pH reversed-phase (HpRP) peptide fractionation

For whole cell proteome samples HpRP fractionation was conducted on an Ultimate 3000 UHPLC system (Thermo Scientific) equipped with a 2.1 mm × 15 cm, 1.7μ Aqcuity BEH C18 column (Waters, UK). Solvent A was 3% ACN, Solvent B was 100% ACN, solvent C was 200 mM ammonium formate (pH 10). Throughout the analysis solvent C was kept at a constant 10%. The flow rate was 400 μL/min and UV was monitored at 280 nm. Samples were loaded in 90% A for 10 min before a gradient elution of 0–10% B over 10 min (curve 3), 10-34% B over 21 min (curve 5), 34-50% B over 5 mins (curve 5) followed by a 10 min wash with 90% B. 15 s (100 μL) fractions were collected throughout the run. Peptide containing fractions were orthogonally recombined into 24 fractions (i.e. fractions 1, 25, 49, 73, 97 combined) and dried in a vacuum centrifuge. Fractions were stored at −80°C prior to analysis.

#### Mass spectrometry

Data were acquired on an Orbitrap Fusion mass spectrometer (Thermo Scientific) coupled to an Ultimate 3000 RSLC nano UHPLC (Thermo Scientific). HpRP fractions were resuspended in 20 μl 5% DMSO 0.5% TFA and 10uL injected. Fractions were loaded at 10 μl/min for 5 min on to an Acclaim PepMap C18 cartridge trap column (300 um × 5 mm, 5 um particle size) in 0.1% TFA. After loading a linear gradient of 3–32% solvent B was used for sample separation over a column of the same stationary phase (75 μm × 50 cm, 2 μm particle size) before washing at 90% B and re-equilibration. Solvents were A: 0.1% FA and B:ACN/0.1% FA. 3h gradients were used for whole cell proteomics samples, 1h gradients for MS-IPs.

An SPS/MS3 acquisition was used for TMT experiments and was run as follows. MS1: Quadrupole isolation, 120’000 resolution, 5e5 AGC target, 50 ms maximum injection time, ions injected for all parallelisable time. MS2: Quadrupole isolation at an isolation width of m/z 0.7, CID fragmentation (NCE 35) with the ion trap scanning out in rapid mode from m/z 120, 8e3 AGC target, 70 ms maximum injection time, ions accumulated for all parallelisable time. In synchronous precursor selection mode the top 10 MS2 ions were selected for HCD fragmentation (65NCE) and scanned out in the orbitrap at 50’000 resolution with an AGC target of 2e4 and a maximum accumulation time of 120 ms, ions were not accumulated for all parallelisable time. The entire MS/MS/MS cycle had a target time of 3 s. Dynamic exclusion was set to +/−10 ppm for 90 s, MS2 fragmentation was trigged on precursor ions 5e3 counts and above. For MS-IPs, MS2 instead used HCD fragmentation (NCE 34) and a maximum injection time of 250ms and had a target cycle time of 2s. For pSILAC MS1 was acquired at 240’000 resolution.

#### Data processing and analysis

For TMT labelled samples data were searched by Mascot within Proteome Discoverer 2.1 in two rounds of searching. First search was against the UniProt Human reference proteome (26/09/17), the HIV proteome and compendium of common contaminants (GPM). The second search took all unmatched spectra from the first search and searched against the human trEMBL database (Uniprot, 26/09/17). The following search parameters were used. MS1 Tol: 10 ppm, MS2 Tol: 0.6 Da. Enzyme: Trypsin (/P). MS3 spectra were used for reporter ion based quantitation with a most confident centroid tolerance of 20 ppm. PSM FDR was calculated using Mascot percolator and was controlled at 0.01% for ‘high’ confidence PSMs and 0.05% for ‘medium’ confidence PSMs. Normalisation was automated and based on total s/n in each channel. Protein/peptide abundance was calculated and output in terms of ‘scaled’ values, where the total s/n across all reporter channels is calculated and a normalised contribution of each channel is output. Proteins/peptides satisfying at least a ‘medium’ FDR confidence were taken forth to statistical analysis in R. This consisted of a moderated T-test (Limma) with Benjamini-Hochberg correction for multiple hypotheses to provide a q value for each comparison (Schwammle et al., 2013). MS-IPs were submitted to a similar search workflow with quantitative data being derived from MS1 spectra via proteome discover minora feature detector node. For pSILAC experiments data were processed in MaxQuant and searched using Andromeda with similar search parameters (Cox and Mann, 2008). MaxQuant output was uploaded into Perseus for calculation of significance B (Tyanova et al., 2016). Where conditions were not carried out in triplicate, downstream analysis was limited to proteins identified with at least 3 unique peptides.

